# Network inference at single-cell resolution reveals morphoregulatory divergence and multilayered buffering in limb evolution

**DOI:** 10.64898/2026.11.07.765081

**Authors:** A Alcaina-Caro, I Sospedra-Arrufat, V Tissières, I Salvador-Martínez, L Almorox, JJ Pawluk, A Díaz-Rueda, A Fernández-Miñán, A Hurtado, FM Real, DG Lupiáñez, B Pernaute, A Visel, M Osterwalder, H Heyn, JJ Tena, G Nusspaumer, J Lopez-Rios

## Abstract

How changes in conserved developmental genetic programs contribute to morphological evolution remains poorly understood, particularly in non-conventional model species. Here, we combine single-cell transcriptomic and chromatin-accessibility profiling with cross-species label transfer and gene regulatory network inference to compare mouse and pig limb development. Our analyses identify broadly conserved cellular organization but discrete divergence in HOX-, FGF- and SHH-regulated programs. FGFR inhibition in mouse limb buds phenocopies the distal expansion of pig *GREM1* expression, while altered SHH responses and anterior-posterior polarity converge on the *PTCH1* regulatory landscape. Its functional dissection reveals degeneration of a limb regulatory module in artiodactyls, conservation of an additional distal enhancer, and extensive redundancy among cis-regulatory inputs. Allele-specific analyses further show that reduced *Ptch1* dosage is buffered by transcriptional compensation. Together, these findings reveal how cis-regulatory and pathway-level robustness can accommodate distributed regulatory change while preserving developmental function, providing a mechanistic framework for the evolution of artiodactyl limb morphology.

## Introduction

The tetrapod limb provides a powerful system for investigating how conserved developmental programs generate morphological diversity. Its homologous segments can be compared across species, the core patterning pathways are unusually well defined, and variation in digit number, skeletal proportions or tissue architecture can often be linked to specific phases of embryonic patterning and growth. Accordingly, comparative studies have uncovered widespread regulatory divergence between homologous limb buds, distinct patterning and post-patterning routes to mammalian digit loss and gene-expression programs associated with differential limb growth^1–5^. Reconstructing the molecular basis of morphological evolution remains nevertheless intrinsically difficult. The causal developmental changes occurred in ancestors that cannot be examined experimentally, while extant lineages separated by millions of years differ at innumerable coding and non-coding sites. Observed molecular differences may therefore represent causes, consequences or compensatory responses which usually cannot be experimentally addressed in non-genetic models. Classical evo-devo emphasized changes in the cis-regulatory regions of conserved developmental genes as a route around pleiotropy^6^, but growing evidence indicates that signalling pathways and transcription factors that pattern embryonic form are particularly robust at the cis-regulatory level^7–11^. Hence, solid morphoregulatory architectures are required to ensure developmental robustness, while simultaneously allowing partial, quantitative or spatially/temporally restricted changes to accumulate during evolution without compromising organismal viability.

Artiodactyl limb evolution provides a particularly informative model to study how developmental robustness can accommodate morphological change. In terrestrial artiodactyls, the weight-bearing axis of the autopod is displaced between the enlarged central digits III and IV, while the flanking digits are reduced or lost. Comparative embryological analyses have shown that similar patterns of digit reduction can arise through distinct developmental routes, involving alterations in early patterning or later tissue remodelling^1,2,12^. In cattle, the acquisition of a symmetrical autopod and digit loss/reduction is accompanied by a progressive loss of anterior-posterior (AP) molecular polarity. Although the posterior SHH source is maintained, graded mesenchymal upregulation of PTCH1 is attenuated and its LRM enhancer has functionally degenerated^1^. Pig limb buds likewise display altered anterior-posterior patterning, including changes in the spatial distribution of *PTCH1* and other HH-responsive genes^2,3^. Together, these observations support a model in which evolution of the symmetric artiodactyl autopod involved selective attenuation of PTCH1-mediated ligand-dependent antagonism.

PTCH1 occupies an unusual position within the HH pathway because it antagonizes signalling through two mechanistically separable activities^13^. In the absence of the ligand, basal PTCH1 represses SMO and prevents pathway activation, a function termed ligand-independent antagonism (LIA)^14,15^. HH binding relieves this repression and activates GLI transcription factors, which induce *Ptch1* among other target genes^16–18^. The resulting accumulation of ligand-binding PTCH1 sequesters SHH and restricts the magnitude and spatial range of signalling, establishing the negative feedback termed ligand-dependent antagonism (LDA)^13,15^. Genetic analyses in *Drosophila* and in the vertebrate neural tube have uncoupled these two activities. HH-insensitive forms of Patched retain cell-autonomous pathway repression but fail to constrain ligand spread^19^, while low constitutive PTCH1 expression preserves LIA even when inducible feedback is removed^13,15^, including the collective feedback mediated by PTCH1, PTCH2 and HHIP^20^. Quantitative changes in the inducible *Ptch1* regulatory landscape could therefore alter HH distribution and tissue patterning while preserving the basal pathway restraint. How such selective changes are accommodated within the broader and potentially redundant *Ptch1* cis-regulatory landscape^21^ remains unknown.

Addressing these questions requires evolutionary differences to be resolved within homologous embryonic cell states and candidate mechanisms to be tested experimentally. Recent advances in single-cell transcriptomic and chromatin-accessibility profiling now allow extensive molecular information to be recovered from the limited embryonic material available for non-model mammals, allowing to identify divergence in gene regulatory network topology or reveal the evolutionary redeployment of conserved developmental programs, as recently shown for the bat wing^5^. Here, we generated matched single-cell transcriptomic and chromatin-accessibility maps of mouse and pig forelimb buds and reconstructed cell state-specific gene regulatory networks. This analysis uncovered discrete divergence in otherwise conserved HOX-, FGF- and HH-associated programs. Guided by previous evidence linking altered HH signalling to the evolution of the artiodactyl autopod, we focused our mechanistic analysis on *Ptch1*. Through cross-species enhancer assays and extensive genetic dissection of the mouse *Ptch1* locus, we show that its limb expression is buffered by distributed enhancer activity and that reduced *Ptch1* dosage is accompanied by increased transcriptional output from the remaining wild-type allele, revealing how an essential developmental locus can remain robust while its transcriptional output evolves.

## Results

### Single-cell multiomic profiling of mouse and pig forelimb buds

Mice have retained the prototypic pentadactyl configuration of ancestral mammals, adapted to their plantigrade/semi-digitigrade locomotion. In contrast, and as all extant terrestrial artiodactyls, pigs have lost the most anterior digit and markedly reduced the lateral ones, yielding a symmetric autopod in which the robust central digits III and IV bear most of the animal’s weight in an unguligrade stance. This digit configuration is accompanied by additional specializations, including loss of the clavicle, powerful girdle elements (scapula and pelvis) and elongated autopod bones, all of which represent biomechanical adaptations associated with cursorial locomotion in these large mammals (Fig. 1a). To build a high-resolution map allowing us to investigate the genetic mechanisms underlying the divergent morphologies of the pentadactyl mouse and the artiodactyl pig limb skeletons, we generated linked single-cell transcriptomic and chromatin accessibility profiles from equivalent forelimb bud stages in the mouse (E10.5/E11.5) and pig (D21/D23)^3^. Joint embedding and unsupervised clustering of the mouse (Table S1) and pig (Table S2) scRNA-seq data recovered the expected cellular compartments of developing limb buds for both species, including mesenchymal progenitors (*Prrx1*^+^/*PRRX1*^+^), ectodermal populations (*Trp63*^+^/*TP63*^+^), myogenic progenitors (*Pax3*^+^/*PAX3*^+^), Schwann cell precursors (*Cdh19*^+^/*CDH19*^+^), endothelial cells (*Pecam1*^+^/*PECAM1*^+^), leukocytes (*Ptprc*^+^/*PTPRC*^+^) and erythroid cells (*Hba-a1*^+^/*HBA-A1*^+^) (Fig. S1). Consistent with tissue dissociation differences between species (see Methods), the single-cell pig datasets showed reduced recovery of ectodermal cells relative to mouse, as evidenced by the reduced number of *TP63*^+^ cells (Fig. S1, arrow) and undetectable *FGF8* expression, despite its previously documented expression in the pig AER^3^, while the mouse ortholog was robustly detected in AER-derived mouse forelimb cells (Fig. S1). Otherwise, all mesenchymal and associated non-ectodermal compartments were represented in both species.

**Figure 1.**
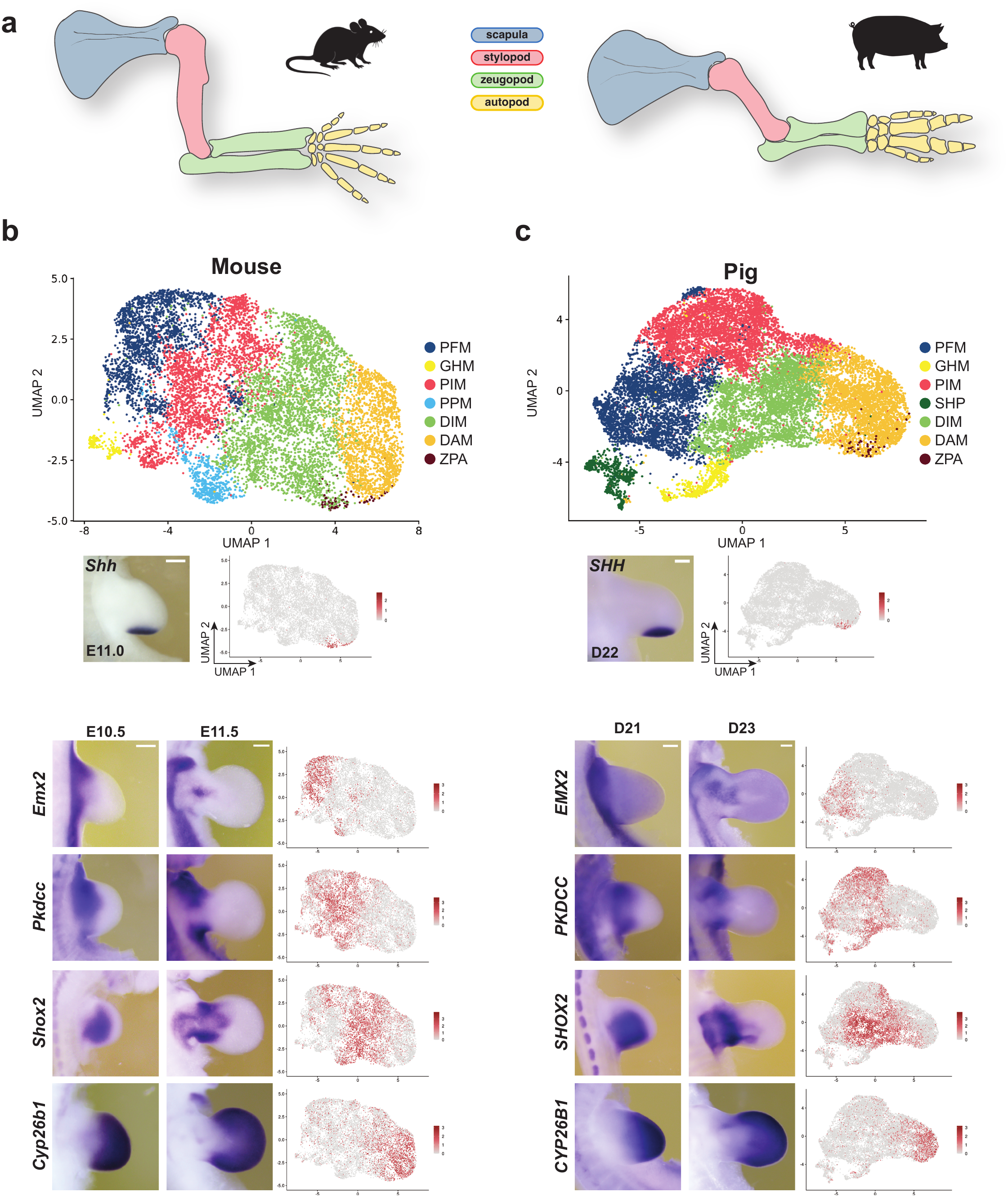
The single-cell landscape of mouse and pig forelimb bud mesenchyme. (a) Schemes of mouse and pig fetal forelimb skeletons. (b, c) UMAP visualization of the mouse (b) and pig (c) forelimb bud mesenchyme scRNA-seq datasets with annotated clusters. Whole-mount in situ hybridization (WISH) analyses for representative cluster-marker genes (*Shh/SHH, Emx2/EMX2*, *Pkdcc/PKDCC*, *Shox2/SHOX2* and *Cyp26b1/CYP26B1*) in E10.5/E11.0/E11.5 (mouse; n=3) and D21/D22/D23 (pig; n=2) forelimb buds are shown next to feature plots depicting the corresponding transcript distributions across the UMAP embedding. PFM: proximal flank mesenchyme; GHM: *Gata4/6*-high mesenchyme; PIM: proximal intermediate mesenchyme; PPM: proximal posterior mesenchyme; SHP: *SIM1*-high population; DIM: distal intermediate mesenchyme; DAM: distal autopod mesenchyme; ZPA: zone of polarizing activity. Scale bar: 0.25mm

We focused our subsequent analyses on the limb mesenchymal compartment in both species, which contains the skeletal progenitor cells, and aimed to define broad populations that helped in cross-species comparisons (Fig. 1b,c). In the mouse, unsupervised subclustering of the forelimb mesenchyme resolved six transcriptionally distinct mesenchymal populations, which we termed PFM (proximal flank mesenchyme); GHM (*Gata4/6*-high mesenchyme); PIM (proximal intermediate mesenchyme); PPM (proximal posterior mesenchyme); DIM (distal intermediate mesenchyme) and DAM (distal autopod mesenchyme). These clusters were supported by characteristic marker combinations, with their relative position within the UMAP projection strongly influenced by proximo-distal transcriptional signatures, as previously noted^22–24^. Four of the clusters (PFM, GHM, PIM and PPM) had evident proximal identity, as indicated by *Meis1*/*Meis2* expression (Fig. S1) and were characterized by the relative enriched expression of specific markers such as *Emx2* (PFM), *Gata4/6* (GHM), *Pkdcc* (PIM) or *Osr1* (PPM). DIM and DAM represented cell populations distal to the previous clusters, with most of the *Shox2*^+^ cells mapping to the DIM population, while DAM expressed typical autopod markers such as *Cyp26b1* or *Tfap2b*. Based on *Shh* expression, we extracted the ZPA population, which was originally included into the DAM cluster (Fig. 1b and Fig. S1). In pig, we also obtained six mesenchymal clusters, five with clear marker-based correspondence to mouse populations (PFM, GHM, PIM, DIM, DAM). A cluster equivalent to mouse PPM was not recovered, and instead we identified an additional cluster with proximal characteristics (SHP: *SIM1*-high population) (Fig. 1c; Fig. S1b). To connect transcriptional states to their anatomical context, we validated the distribution of cluster-enriched markers by whole-mount in situ hybridization (WISH) in mouse (E10.5/E11.5) and pig (D21/D23) limb buds. These analyses confirmed the spatially restricted expression patterns for pig *EMX2*, *PKDCC*, *SHOX2*, *CYP26B1* and *SHH*, which were detected in similar domains as their well-known mouse orthologues (Fig. 1b and 1c).

### Cross-species label transfer enables comparison of mesenchymal states in a shared transcriptomic space and identifies distal FGF signalling divergence in pig limb buds

Next, to compare transcriptional profiles across corresponding mouse and pig forelimb mesenchymal states, we used a label transfer strategy using ortholog genes to map pig mesenchymal cells onto the annotated mouse reference atlas and project them into its UMAP embedding (Figs. 2a,b and S2a-c). As expected, pig mesenchymal populations aligned well with the equivalent populations in the mouse, showing a high correspondence across the conserved populations (PFM, GHM, PIM, DIM, DAM and ZPA; Fig. 2c). This mapping also indicated the absence of a clear pig PPM equivalent population, as pig cells mapping onto the mouse PPM coordinates were of mixed origin. Conversely, the pig SHP population was mostly distributed across the mouse proximal clusters PFM and GHM (Fig. 2c). Feature plots for the same marker genes used in Fig. 1 (*EMX2/Emx2* for PFM, *PKDCC/Pkdcc* for PIM, *SHOX2/Shox2* for DIM, *CYP26B1/Cyp26b1* for DAM and *SHH/Shh* for ZPA) showed a clear match between the mouse and pig-on-mouse UMAPs, indicating that the defining transcriptional signatures for each of the clusters remain coherent after cross-species projection (Fig. S2d). WISH for markers (*Wt1*/*WT1*, *Gsc*/*GSC*, *Pax1*/*PAX1*, *Osr1*/*OSR1*, *Osr2*/*OSR2*, *Tbx2*/*TBX2 and Hoxd11*/*HOXD11*) (Fig. S2e) validated that our label transfer strategy constitutes a valuable tool to compare gene expression profiles and homologous mesenchymal states between the two species within the same coordinate system.

**Figure 2.**
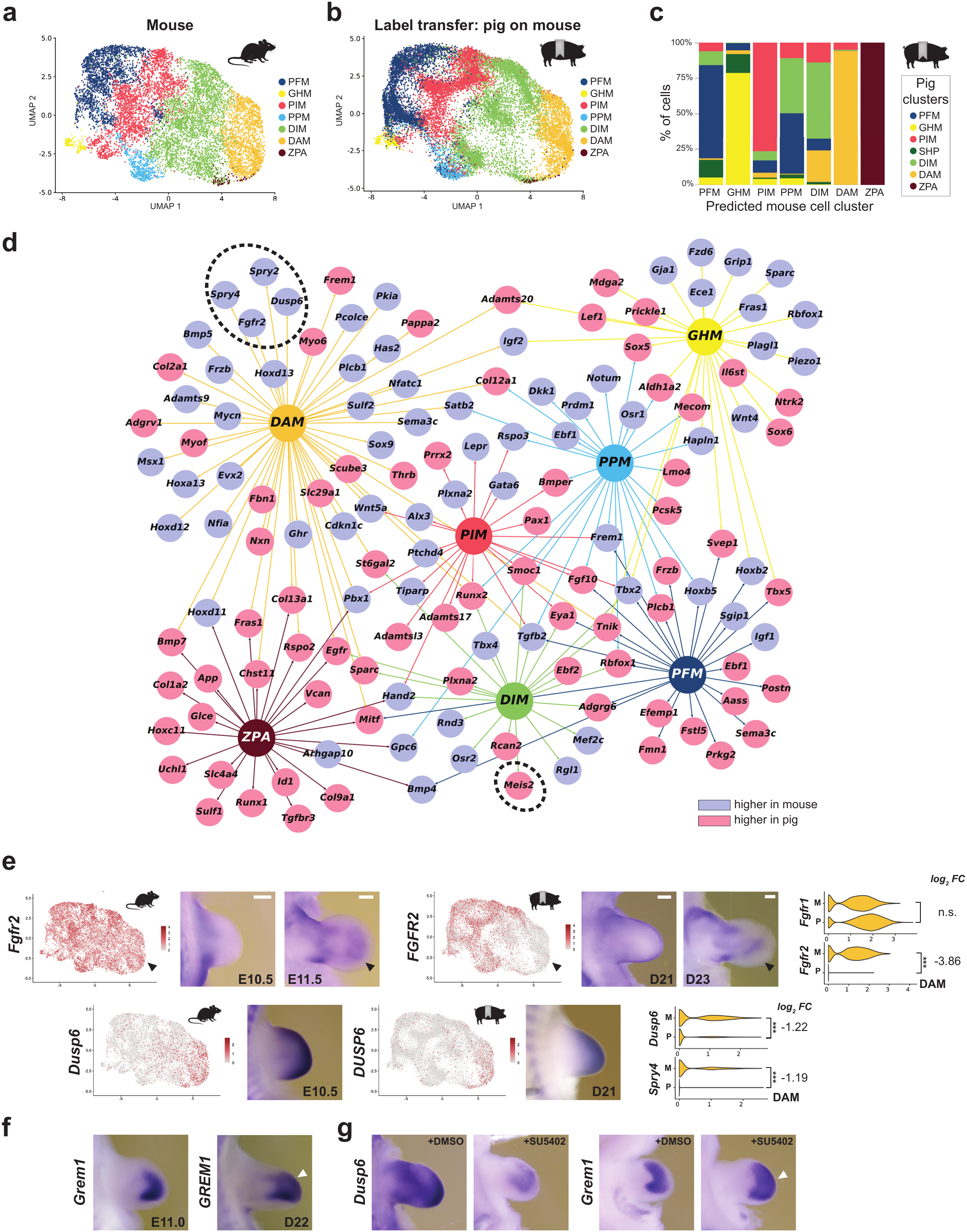
Label transfer integration of mouse and pig limb bud mesenchymal scRNA-seq datasets reveals altered FGF signalling in the distal pig limb bud mesenchyme. (a) UMAP representation of the mouse forelimb mesenchyme scRNA-seq dataset with annotated populations (same as Fig. 1b). (b) UMAP representation of the cross-species label transfer from the pig to the mouse dataset. (c) Bar plot displaying the proportions of native pig cluster labels within each mouse-predicted cluster. (d) Network plot of differentially expressed genes associated with limb development (LP: limb phenotype, Tables S3 and S4), curated from Gene Ontology and Mammalian Phenotype categories related to limb morphogenesis. Each LP gene is assigned to the mesenchymal domain in which it is differentially expressed between mouse and pig, with node color indicating the direction of change in pig versus mouse cells (pink, upregulated in pig; blue, upregulated in mouse). The upregulation of *MEIS2* in DIM cells and the downregulation of *FGFR2* and the FGF pathway transcriptional readouts *DUSP6* and *SPRY2/4* are indicated with dashed circles. (e) Expression of *Fgfr2/FGFR2* and *Dusp6/DUSP6* on the mouse and pig UMAP projections (left) and corresponding WISH at equivalent developmental stages (mouse E10.5/E11.5; pig D21/D23). Violin plots (right; also shown for *Fgfr1*/*FGFR1*) represent normalized expression distributions across individual cells, with adjacent values indicating the pig-versus-mouse log_2_ fold change (FC) in the DAM population at the cell level; ***p ≤ 0.001). (f) *Grem1/GREM1* transcript distribution in mouse (E11.0) and pig (D22) limb buds. The characteristic *Grem1*-free region in the sub-AER mesenchyme of mouse limb buds is absent in pig limb buds, where the *GREM1* expression domain becomes distalized (white arrowheads). (g) E10.5 mouse forelimbs cultured for 16 hours in the presence of the FGFR-inhibitor SU5402 display downregulation of *Dusp6* and distal expansion of *Grem1* (white arrowhead) compared with DMSO-treated contralateral controls. n=3 for all mouse and n=2 for all pig WISH. Scale bar: 0.25mm.

In addition to providing comparable visual representations of individual gene expression profiles at the single cell level, our label transfer strategy also allowed us to perform exploratory differential gene expression analyses between equivalent mouse mesenchymal populations and their mapped pig counterparts, thus enabling us to identify genes whose expression in the limb bud has diverged since their last common ancestor (Table S3). To exclude ubiquitously expressed genes, we restricted the analysis to genes that were differentially expressed between clusters in at least one of the two species (see Methods). The number of candidate differentially expressed genes (DEG) was 170 in PFM, 213 in GHM, 107 in PIM, 141 in PPM, 99 in DIM, 309 in DAM and 126 in ZPA (adjusted p<0.05; absolute average log_2_ Fold change >1). Gene ontology (GO) analysis for biological processes on these DEG sets showed enrichment in skeletal/limb morphogenesis terms in all clusters, with the exception of DIM (Fig. S3). In addition, and to identify specific genes whose expression divergence may have contributed to anatomical limb diversification in pigs, we retrieved the list of genes with known functions in limb development in mammals (LP: limb phenotype; Table S4) and inquired which of them were differentially expressed between mouse and pig limb bud equivalent clusters. Network representation of these LP DEG allowed for the visualization of the most prominently down- or upregulated genes in pig versus mouse limb buds (in blue and pink, respectively) for each of the mesenchymal subpopulations (Fig. 2d). While differences in LP gene expression are broad across all cell clusters, and consistent with the previous analyses, the DAM cluster contained the largest number of genes meeting these differential-expression thresholds (Table S3). These observations are relevant, as the handplate is the limb skeletal segment that has most profoundly deviated from the archetypical tetrapod blueprint that has been retained in the mouse. Quite strikingly, this analysis revealed a downregulation of the *FGFR2* gene in pig handplate progenitor cells, simultaneous with diminished expression of robust FGF signalling transcriptional readouts such as *DUSP6, SPRY2 and SPRY4*^25^. Concomitantly, we observed an upregulation of the proximal marker *MEIS2* in the DIM population (Fig. 2d). While *MEIS2* upregulation in pig limb buds is evident at the single cell level, the resolution provided by WISH analysis was insufficient (Fig. S4). In contrast, we detected very clear differences in the expression of *Fgfr2*/*FGFR2* between mouse and pig limb buds, as *FGFR2* expression domain appeared markedly reduced in pig in comparison to its orthologue expression in mouse handplates (arrowheads in Fig. 2e), a difference clearly seen as well in the corresponding UMAPs. In line with putatively reduced FGF signaling in DAM cells, we confirmed by WISH the downregulation of *DUSP6* expression in pig limb buds (Fig. 2e). These results suggest that the pig distal-most limb mesenchymal cells display a diminished activation of the FGF pathway. Previous genetic studies in the mouse revealed a requirement for FGFR activity in preventing *Grem1* expression in the posterior sub-ectodermal limb bud mesenchyme^26^. As the distal expansion of an almost symmetric *Grem1* domain is a shared landmark of artiodactyl limb buds^1,3^ (Fig. 2f), we inquired if treatment of mouse limb buds with the FGF receptor inhibitor SU5402 would suffice in triggering the distal expansion of *Grem1* expression. Indeed, culturing mouse limb buds in the presence of 10µM SU5402 elicited a strong downregulation of *Dusp6* and the loss of the characteristic gap devoid of *Grem1* expression in the distal handplate, which is reminiscent of *GREM1* expression in pig limb buds (Fig. 2g).

### Conserved limb regulatory architecture and species-specific divergence of HOXB and HOXC programs

Having identified candidate cross-species transcriptional differences by differential gene expression analysis, we next applied SCENIC+^27^ to the linked single cell transcriptomic and chromatin accessibility profiles generated from mouse and pig limb buds. SCENIC+ infers enhancer-driven regulons (eRegulons) by linking upstream transcription factors to candidate target genes through accessible regulatory regions containing compatible transcription factor binding motifs, and integrating transcription factor expression, target gene expression and chromatin accessibility within a unified regulatory framework. We performed this analysis using both direct (Fig. 3) and extended (Fig. S5) motif annotations (see Methods). This analysis revealed a substantial shared regulatory architecture consistent with the spatial and differentiation identities of the annotated mouse and pig limb bud populations (Fig. 3a,b and Fig. S5a,b; see Tables S5 and S6 for ranked top-40 regulons per cluster). For instance, proximal mesenchymal populations were characterized, among others, by MEIS1/PBX1/NR2F1/NR2F2/GATA4/GATA6-associated programs, distal populations by HOXA13/HOXD13/TFAP2B/LHX2/LHX9/MSX1-associated regulons, and differentiating skeletal populations by SOX9/RUNX2-associated activities. These results indicate that the inferred regulon landscape largely recapitulates the expected regulatory logic of the developing limb bud. We also validated the ability of SCENIC+ to recover long-range regulatory interactions within complex cis-regulatory landscapes. Indeed, SCENIC+ successfully captured previously characterized limb enhancers distributed over large genomic intervals surrounding *Grem1*, *Shox2* and *Gli3*, whose regulatory landscapes have been extensively dissected at the functional level^7,9,10,28^ (Fig. S6 and Table S7).

**Figure 3.**
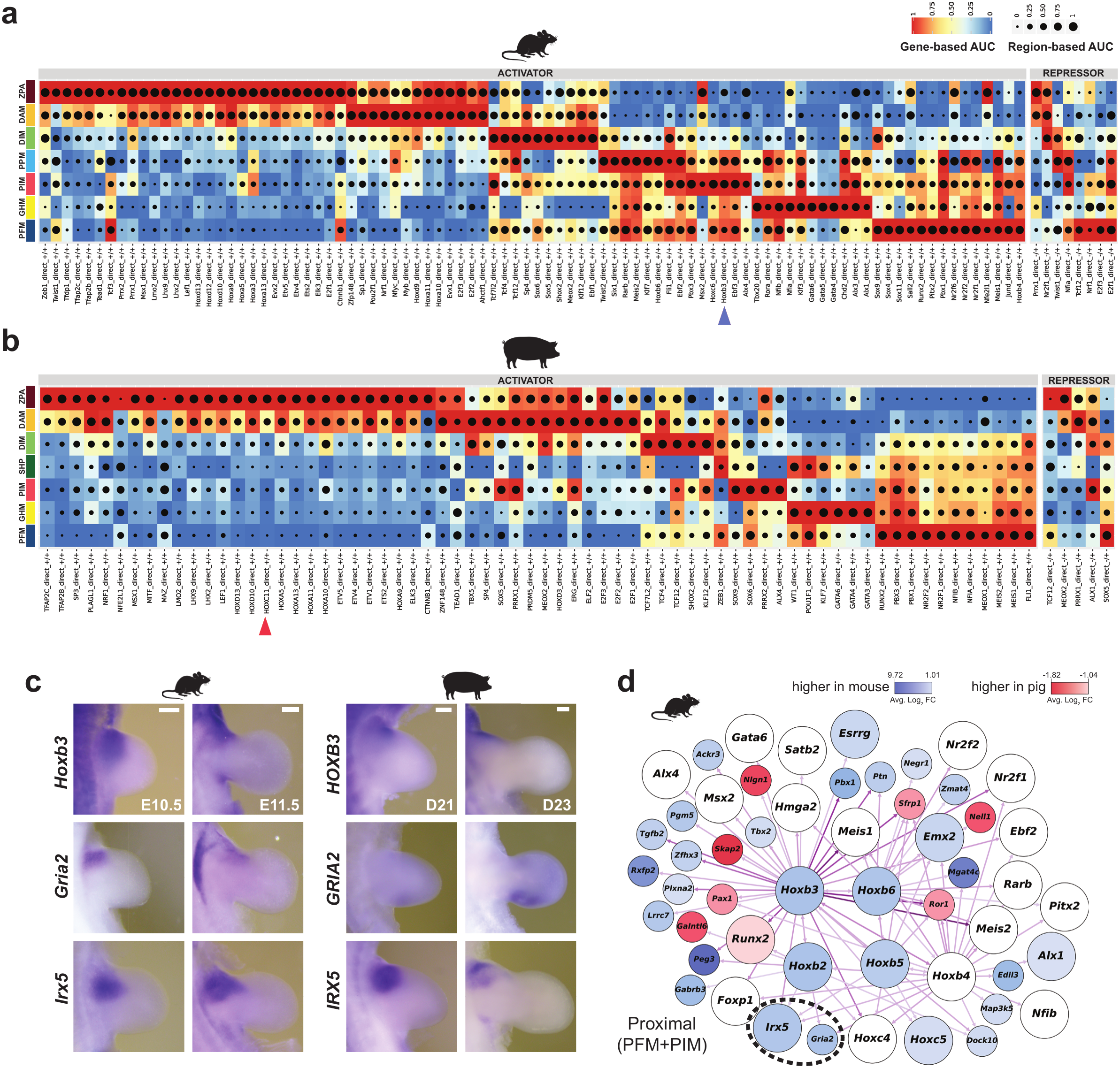
Conserved and divergent regulatory programs in mouse and pig limb buds. (a, b) SCENIC+ heatmaps of the mouse (a) and pig (b) single-cell multiomic datasets for activator and repressor transcription factor (TF) networks. Columns indicate the TF with predicted direct target relationships (regulons), with rows representing mesenchymal clusters. The heatmap and dot size reflect gene-based and region-based mean AUC scores, respectively. A blue arrowhead indicates *Hoxb3* regulon activity in mouse proximal clusters, which is not detected by SCENIC+ in pig datasets. Red arrowhead indicates the species-specific activation of the *Hoxc11* regulon in pig DAM and ZPA populations. See also Fig. S5. (c) WISH validation of the differential expression of *Hoxb3/HOXB3*, *Gria2/GRIA2* and *Irx5/IRX5* at equivalent developmental stages in mouse (E10.5/E11.5; n=3 per stage) and pig (D21/D23; n=2 per stage), suggesting species-specific rewiring of proximal TF networks. (d) Network representation of the mouse *Hoxb2*-*Hoxb6* targets in the main proximal clusters (PFM and PIM) as predicted by SCENIC+. Large nodes correspond to transcription factors belonging to the top 20 transcription factors within the proximal clusters. Small nodes denote predicted *Hoxb2*-*6* transcriptional targets up- or downregulated in pig in proximal cells, labelled pink and blue, respectively. Only targets that passed the interspecies differential expression filters or ranked among the top 20 TFs by RSS in PFM/PIM are shown. Arrow intensity reflects the correlation between TF expression and expression of its predicted target (TF2G rho). Scale bar: 0.25mm.

Our analysis using SCENIC+ also revealed marked species-specific regulatory signatures. Most notably, while proximal mouse mesenchymal clusters were enriched for HOXB-associated regulons, including HOXB2-HOXB6 (blue arrowheads in Fig. 3a,b and Fig. S5a,b), these transcriptional programs were not detected in the corresponding pig populations (Tables S5 and S6). Consistent with this regulatory divergence, differential expression analysis showed that several of these pig *HOXB* genes are expressed at substantially lower levels in proximal pig limb bud populations (Table S3), a difference that we confirmed for *Hoxb3*/*HOXB3* by WISH (Fig. 3c). We next used the SCENIC+ regulon architecture (Table S7) to reconstruct a simplified HOXB candidate target network associated with the proximal mouse PFM and PIM populations, onto which we overlaid cross-species differential expression information (Fig. 3d). This approach identified several putative transcriptional targets expressed at lower (blue) or higher (red) levels in pig limb buds, including *Gria2*/*GRIA2* and *Irx5*/*IRX5*. WISH analysis revealed that *GRIA2* is barely detectable in proximal pig limb buds at either D21 or D23, but is prominently expressed in posterior limb-bud mesenchyme, indicating a marked alteration of its expression pattern relative to the mouse ortholog (Fig. 3c). *Irx5*/*IRX5* expression appeared broadly comparable between species at E10.5/D21, but showed reduced proximal expression in pig D23 limb buds compared with mouse E11.5 limb buds. Interestingly, *Tbx2*, predicted by SCENIC+ as a HOXB3 target, showed reduced expression in the proximal flank mesenchyme of pig limb buds (Fig. S2).

In addition to these proximal alterations, SCENIC+ identified a prominent pig-specific HOXC11 regulon in distal limb bud populations, including the ZPA and DAM clusters (red arrowheads in Fig. 3b and Fig. S5b; Table S6). Reconstruction of the HOXC11-associated network in ZPA cells (Table S8) retrieved a small set of candidate transcriptional targets, several of which were upregulated in pig relative to mouse ZPA cells (Fig. S5c). This observation is particularly striking because *Hoxc11* is expressed in the posterior mesenchyme of mouse and chick hindlimb buds but transcriptionally silent in the forelimb^29,30^, suggesting a redeployment of a hindlimb-associated HOX program in the pig forelimb. Of note, *HHIP*, which encodes a ligand-binding antagonist of Hedgehog signalling, was included among the predicted targets of the pig HOXC11 regulon, suggesting that the HOXC11-associated regulatory divergence may intersect with the HH pathway. Together, these results identify an overall conserved limb regulatory architecture as well as pig-specific attenuation and redeployment of HOX-associated regulatory programs in defined limb bud populations. Moreover, this integrative approach makes the availability of cell-matched scRNAseq and scATACseq datasets a powerful tool to reconstruct gene regulatory networks *de novo*, or with limited *a priori* knowledge, to uncover evolutionary divergence in morpho-regulatory systems.

### Spatial remodeling of Hedgehog responses in pig limbs and functional dissection of the *Ptch1* LRM enhancer

Our analyses so far show that the conserved mesenchymal cell repertoire displays cross-species differences in signalling and regulatory programs, including FGF-responsive and HOX-associated networks. Notably, several of the transcriptional and regulatory differences identified by the cross-species analyses converged on distal mesenchymal populations, which contain the handplate progenitors, a limb segment that has diverged extensively between mouse and pig. A defining feature of the artiodactyl limb is that the anatomical loss of asymmetry in the handplate is preceded by molecular alterations in AP polarity of the distal limb bud mesenchyme. These changes manifest in the artiodactyl limb bud as the distalization and anterior expansion of molecular markers such as *GREM1* or 5’*HOXD* genes^1,3^, together with diminished *LHX2* expression in the anterior mesenchyme^24^. To determine whether this altered polarity was also apparent at single-cell transcriptomic resolution, we performed unsupervised reclustering of the mouse and pig DAM cells. This analysis recovered two populations corresponding to anterior and posterior distal limb bud identities (Fig. 4a). Projection of the two pig DAM subclusters onto the mouse UMAP embedding following label transfer revealed a shift towards posterior-like transcriptional identities in the pig distal mesenchyme (Fig. 4b,c).

**Figure 4.**
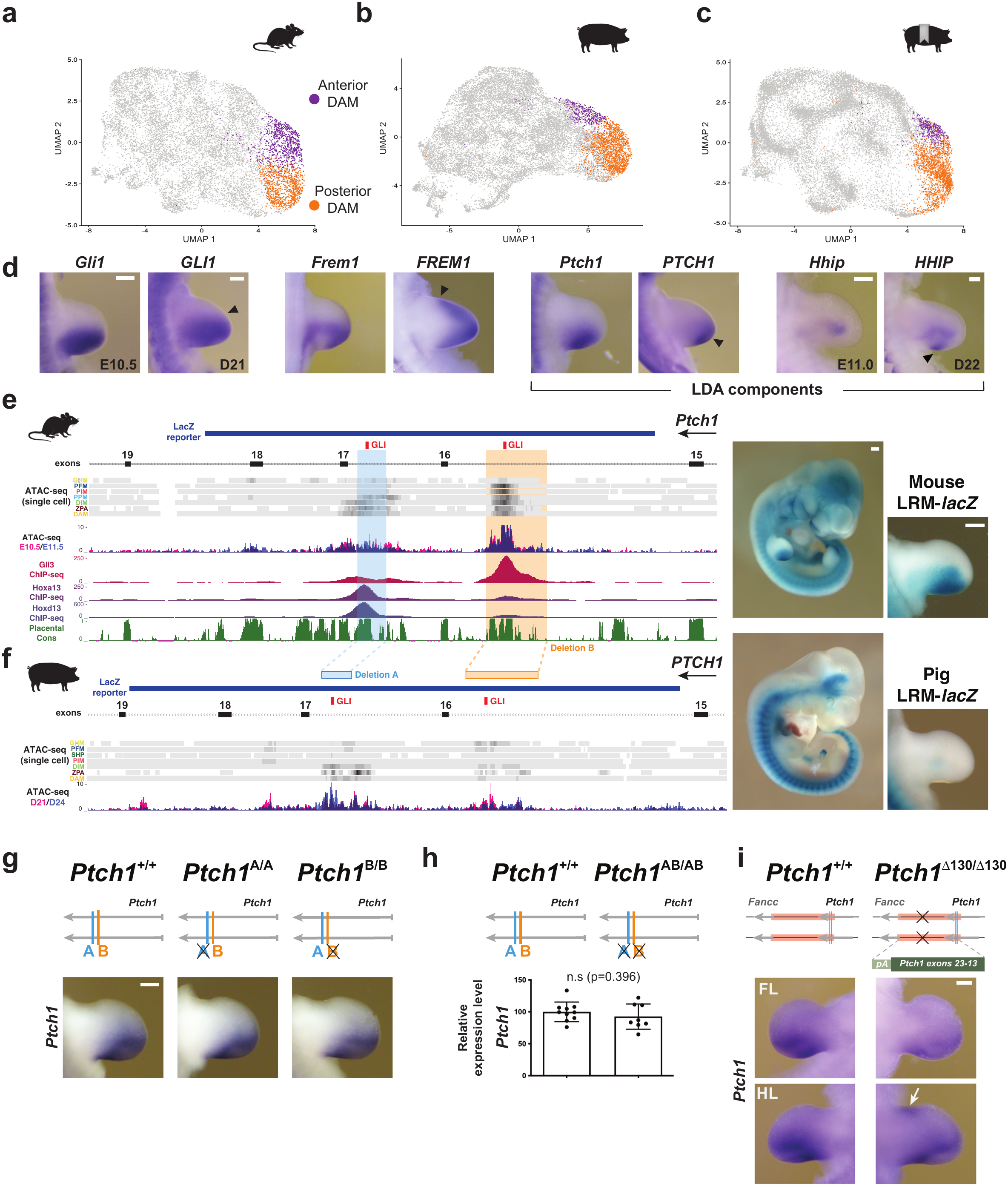
Divergent limb regulatory activity in the *Ptch1* locus correlates with altered anterior-posterior patterning and SHH response in the distal mesenchyme of pig limb buds. (a-c) UMAP representations of mouse (a), pig (b), and pig-on-mouse label transfer (c) datasets indicating anterior (purple) and posterior (orange) DAM subclusters. Label-transfer mapping reveals a shift towards posterior-like transcriptional identities in the pig distal autopod mesenchyme. (d) WISH showing *Gli1/GLI1*, *Frem1/FREM1, Ptch1/PTCH1* and *Hhip*/*HHIP* expression in mouse (n=3) and pig (n=2) limb buds. Pig limb buds display distalized *PTCH1* expression together with an anterior expansion of the HH targets *GLI1* and *FREM1* relative to mouse limb buds, while *HHIP* displays an additional posterior domain (arrowheads). (e) The mouse *Ptch1* LRM enhancer comprises two major regions of open chromatin (denoted A and B) enriched in GLI3 and HOXA13/HOXD13 containing complexes. The mouse LRM drives posterior expression of a *lacZ* reporter in mouse embryonic limbs^1^ (right). Single-cell and bulk ATAC-seq tracks and HOXA13, HOXD13 and GLI3 ChIP-seq profiles in mouse limb buds are shown^3,41,42^, together with the sequence conservation track for placental mammals. (f) The porcine *PTCH1* LRM region exhibits reduced chromatin accessibility in pig limb buds, consistent with diminished reporter activity in limb buds in mouse embryos carrying a pig *LRM-lacZ* reporter transgene (n=6; see Fig.S7a). (g) Whole-mount ISH for *Ptch1* in mouse limb buds from homozygous mutants carrying deletions in either region of open chromatin (A or B) within the LRM (*Ptch1*^A/A^; *Ptch1*^B/B^). A and B deletions are shaded blue and orange in panel (e). Neither deletion alone is sufficient to alter *Ptch1* expression in limb buds when compared to wild-type controls (n=3 per genotype). (h) Quantitation of *Ptch1* expression levels in *Ptch1*^AB/AB^ E11.0 forelimb buds relative to wild-type littermates by RT-qPCR. Data are shown as mean ± SD (n= 8-10 independent embryos per genotype, represented as dots). (i) Mouse embryos homozygous for the *Ptch1*^Δ130/Δ130^ replacement allele show reduced *Ptch1* expression in forelimb and hindlimb buds and ectopic *Ptch1* activation in the anterior margin of hindlimb buds (white arrow) by WISH analysis (n=3). The *Ptch1*^Δ130^ allele replaces 130 Kb, including the entire LRM and all *Ptch1* downstream sequences up to the *Fancc* promoter and exon 2, by a synthetic cassette restoring the *Ptch1* exon 13-23 coding sequence. Scale bar: 0.25mm.

Previous studies by others and us^1–3^ have associated these patterning alterations in artiodactyls with changes in Sonic Hedgehog (SHH) signaling, which plays a central role in AP polarization and growth of the distal limb^31–35^. Although *Shh*/*SHH* expression itself is largely conserved (Fig. 1b), both pig and bovine limb buds display reduced posterior *GLI1* expression together with an anterior shift relative to the mouse pattern^1–3^ (Fig. 4d). Consistent with this altered response, additional HH target genes, including *GREM1*^10,36,37^ (Fig. 2) and *FREM1*^38^ (Fig. 4d), also show anterior shifts. These alterations are accompanied by a distally restricted *PTCH1* expression domain at early stages of pig limb development in comparison to mouse^2,3^ (Fig. 4d). As ligand-dependent upregulation of PTCH1 contributes to the ligand-dependent antagonism mechanism that restricts SHH distribution to the posterior limb mesenchyme^13,15,19,20^, altering the spatial distribution of PTCH1 could modify the range of HH transcriptional target activation in the pig handplate, as previously shown in bovines. Interestingly, analysis of *Hhip*/*HHIP* expression, another component of the ligand-dependent antagonism machinery^20,39^, revealed a largely conserved expression in the core mesenchyme together with an additional posterior domain in pig limb buds that overlaps with the ZPA (Fig. 4d, arrowhead). Overall, these observations indicate that the spatial organization of Hedgehog responses and the expression of genes encoding ligand-dependent antagonism components have been remodeled in the pig limb.

The divergence we detect in the activity and spatial organization of conserved developmental programs raises the question of how regulatory changes can accumulate without compromising essential gene functions. Among these HH pathway components, we focused our mechanistic analyses on *Ptch1* as previous work provided evidence for its key functions in limb AP patterning^40^ and identified a limb regulatory module (LRM) within the *Ptch1* locus that has functionally degenerated in the bovine genome^1^. We therefore asked whether this evolutionary alteration was shared with pigs and how the LRM contributes to *Ptch1* regulation *in vivo*. The mouse LRM is located within the intronic-exonic region of the *Ptch1* locus and contains two prominent regions of accessible chromatin that are bound in limb bud cells by GLI3 and by HOXA13- and HOXD13-containing complexes^41,42^ (Fig. 4e). The orthologous pig LRM displays detectable chromatin accessibility, although the open regions are less prominent than in mouse (Fig. 4f). To assess the regulatory activity of the porcine element, we cloned the pig LRM upstream of a *lacZ* reporter and generated transgenic mouse embryos. Similar to the mouse LRM, the pig element retained reporter activity along the primary body axis. In contrast, its activity in the limb mesenchyme was greatly reduced (n=2/6) or abolished (n=4/6; Fig. 4e,f; Fig. S7a), closely resembling the loss of limb activity previously observed for the bovine enhancer in transgenic reporter assays^1^ (Fig. S7a). These findings indicate that degeneration of the limb enhancer activity of the *PTCH1* LRM is shared by at least two early-diverging artiodactyl lineages and is therefore unlikely to represent a particularity of bovids.

Consistent with a role for the LRM in the ligand-dependent antagonism mechanism that transcriptionally upregulates *Ptch1* in response to HH signaling^16,17^, limb enhancer activity of the LRM is entirely dependent on SHH signaling, as reporter activity was completely abolished in limb buds of *Shh*-deficient embryos (Fig. S7b). Based on this result, and to investigate the contribution of the LRM to *Ptch1* expression in embryonic limbs, we focused initially on the two conserved regions of accessible chromatin bound by GLI-containing complexes (regions A and B; Fig. 4e). We generated targeted deletions encompassing the evolutionarily conserved GLI-binding motifs in either region A or region B individually (*Ptch1*^A^ and *Ptch1*^B^ alleles, respectively), as well as a combined deletion of both regions (*Ptch1*^AB^), without disrupting the splice sites of the adjacent exons. Unexpectedly, homozygous embryos carrying any of these alleles showed no detectable alteration in endogenous *Ptch1* expression in the limb bud (Fig. 4g,h). Similar results were obtained in trans-heterozygous embryos carrying each deletion in combination with a *Ptch1* null allele (Fig. S7c), indicating that removal of these GLI-binding regions alone is insufficient to measurably impair *Ptch1* expression in the limb. Consistent with this observation, a mouse LRM-lacZ reporter lacking both GLI-binding regions (LRM^ΔAB-*lacZ*^) retained residual enhancer activity in the limb mesenchyme (Fig. S7d), indicating that while regions A and B are essential for robust LRM function, additional sequences still mediate some transcriptional activation.

As the LRM is embedded within the coding region of *Ptch1*, complete deletion of the element would simultaneously disrupt gene function. This would preclude a straightforward genetic analysis, as *Ptch1*-deficient embryos die around E9.0, shortly before the limb buds emerge^43^. To overcome this limitation, we used embryonic stem cells (ESCs) to engineer the *Ptch1*^Δ130^ replacement allele in which the ∼130-Kb interval extending from *Ptch1* exon 13 to the adjacent *Fancc* exon 2 was substituted by a synthetic cassette that restores the *Ptch1* exon 13-23 coding sequence while simultaneously eliminating the LRM. Homozygous *Ptch1*^Δ130/Δ130^ ESCs were aggregated with tetraploid embryos to generate entirely ESC-derived embryos^44^. *Ptch1*^Δ130/Δ130^ embryos recovered at E11.5 showed no overt morphological abnormalities, indicating that the synthetic exon 13-23 cassette restores *Ptch1* function and bypasses the early lethality caused by complete *Ptch1* deficiency. WISH analysis revealed a reduction of *Ptch1* expression in the posterior mesenchyme of both fore- and hindlimb buds of *Ptch1*^Δ130/Δ130^ embryos (Fig. 4i). In addition, mutant hindlimb buds displayed an ectopic anterior domain of *Ptch1* expression (Fig. 4i, white arrow), consistent with ectopic anterior activation of *Shh*, as previously reported for the conditional inactivation of *Ptch1* in the limb mesenchyme^40^. These results indicate that the interval containing the LRM contributes to the posterior, SHH-dependent upregulation of *Ptch1* during limb development.

### Multiple distal limb enhancers are distributed across the *Ptch1* regulatory landscape

Transcriptional upregulation of *Ptch1* in limb buds is neither completely abolished in artiodactyls, despite functional degeneration of the LRM, nor in *Ptch1*^Δ130/Δ130^ embryos lacking the endogenous LRM. This indicates that additional cis-regulatory elements contribute to SHH-dependent *Ptch1* upregulation during limb development. In the mouse genome, *Ptch1* is flanked by *Fancc* and *Ercc6l2*, two broadly expressed genes involved in DNA repair (Fig. 5a and Fig.S8a). Both genes are detected at low levels in the limb bud, but their expression domains do not resemble the posterior pattern characteristic of *Ptch1* (Fig. S8b). Previous studies identified several putative *Ptch1* enhancers within the *Ptch1* cis-regulatory landscape (Fig. 5a, blue arrowheads; enhancers [E] renamed with numerals indicating their distance in kilobases from the *Ptch1* transcriptional start site [TSS])^1,18,21,45,46^. Most of these elements display broad activity in neural tissues (Fig. S8c for VISTA-assayed elements), with some also active in other regions, including the gut mesenchyme or the AER^18,21^. Among these previously characterized elements, only the LRM drives robust reporter expression in the limb bud mesenchyme^1^. Nevertheless, GLI3 ChIP-seq from limb buds^41^ revealed additional prominent uncharacterized GLI-bound regions within the *Ptch1* regulatory landscape. These included one region near the *Fancc* promoter and a cluster of sites located more than 300 kb upstream of *Ptch1*, which we termed E335, E351, and E385. To determine if these regions represent candidate limb regulatory elements interacting with the *Ptch1* locus, we performed UMI-4C^47^ using viewpoints centered on the four most prominent GLI-bound regions in the limb bud: the *Ptch1* promoter, the LRM, the *Fancc* promoter, and the distal candidate enhancer E385 (Fig. 5b). This analysis confirmed contacts between the LRM and the *Ptch1* promoter and identified interactions of the *Ptch1* promoter with both the *Fancc* promoter and the distal E385 element. The potential regulatory relevance of these chromatin contacts was independently supported by single-cell multiome peak-to-gene correlation analysis using Signac^48^, which identified long-range associations between *Ptch1* transcription and chromatin accessibility at several regulatory elements, including the LRM and the distal cluster of candidate enhancers (Fig. 5c).

**Figure 5.**
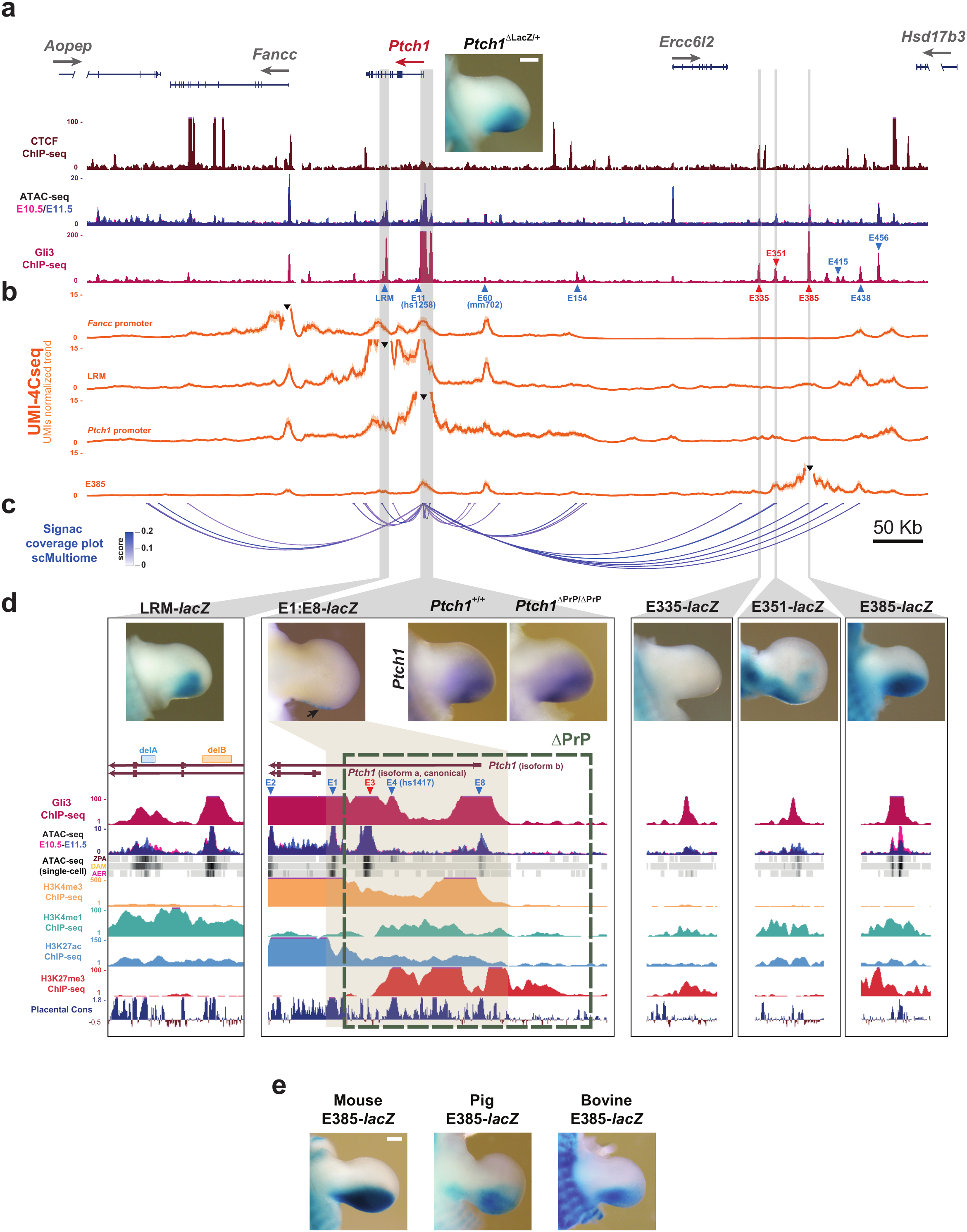
Regulatory architecture and enhancer dissection of the mouse *Ptch1* locus in limb buds. (a) Genomic view of the mouse *Ptch1* cis-regulatory landscape, with the neighboring genes *Aopep*, *Fancc*, *Ercc6l2* and *Hsd17b3* indicated. ATAC-seq^3^, CTCF ^89^ and GLI3 ChIP-seq^41^ profiles from limb buds are also displayed. The inset shows the limb bud expression pattern of the *Ptch1*^ΔlacZ^ reporter allele. Previously identified cis-regulatory regions are indicated with blue arrowheads, including the LRM^1^, VISTA hs1258 and mm702^45,46^ (renamed here as E11 and E60, according to their distance from the canonical TSS), and E154, E415, E438 and E456^21^. Red arrowheads indicate the newly identified accessible regions E335, E351 and E385, which show prominent enrichment in GLI3-containing complexes. (b) UMI-4C-seq interaction profiles across the *Ptch1* cis-regulatory landscape using the *Fancc* promoter, the LRM, the *Ptch1* promoter and the E385 element as viewpoints (black arrowheads; n=2 per viewpoint). (c) Signac correlation links between chromatin accessibility and *Ptch1* expression in mouse limb buds. Color scale represents the Pearson correlation value. (d) Functional dissection of *Ptch1* limb bud cis-regulatory elements. Left: LRM-lacZ reporter and genomic profiling of the LRM enhancer (see Figure 4). Center: the *Ptch1* promoter proximal region contains elements E1, E3, E4 and E8. The TSS of the canonical *Ptch1* isoform (isoform a) and of isoform b are indicated. The reporter activity of the isolated proximal elements was previously tested (E1^1,18^; E4 [hs1417^45,46^]; E2 and E8^21^). None of these regions is able to drive lacZ reporter expression in the limb mesenchyme of transgenic mouse embryos, and neither did the E3 element (Fig. S8d). A 8.7-Kb construct spanning the E1-E8 region (E1:E8-*lacZ*; shaded in grey) drives reporter expression in the limb to the posterior AER (n=2/2), similar to the E1 minimal promoter element alone^1,18^. Although the ΔPrP deletion removes all regions of open chromatin upstream of the minimal promoter E1, *Ptch1* expression in the limb bud is unaltered in homozygous *Ptch1*^ΔPrP/ΔPrP^ embryos compared to wild-type littermates. Right: *lacZ* reporter expression driven by the distal candidate elements E335 (n=2/2), E351 (n=2/2) and E385 (n=3/3) in transgenic mouse limb buds (see also Fig. S8). GLI3 ChIP-seq, bulk and single-cell ATAC-seq for selected clusters, H3K4me3, H3K4me1, H3K27ac, H3K27me3 ChIP-seq profiles and placental conservation tracks are shown for all highlighted regions^41,90^. (e) Mouse forelimbs showing reporter activity driven by the orthologous mouse (n=3/3), pig (n=3/3) and bovine (n=3/5) E385-*lacZ* constructs, revealing conserved enhancer activity in the distal posterior limb bud. Scale bar: 0.25mm.

The high density of GLI-bound regions surrounding the *Ptch1* promoter prompted us to investigate whether promoter-proximal elements might collectively contribute to limb-specific *Ptch1* upregulation. Previous studies showed that none of the individual promoter-associated regions E1, E2, E4, or E8 displayed mesenchymal enhancer activity when tested separately^1,18,21,45,46^. While the remaining untested E3 region was similarly inactive in the limb mesenchyme (Fig. S8d), we nevertheless reasoned that these elements might function cooperatively. To evaluate this possibility, we generated a reporter construct encompassing the complete E1:E8 interval upstream of the *Ptch1* promoter. This extended region failed to drive reporter expression in the limb mesenchyme and instead recapitulated the posterior AER activity previously observed for the proximal E1 promoter element alone^1,18^ (Fig. 5d). Consistent with these reporter assays, deletion of the complete upstream interval in the *Ptch1*^ΔPrP^ allele, which left the minimal E1 promoter region untouched, did not alter *Ptch1* expression in the posterior limb mesenchyme, despite removing the TSS of the alternative *Ptch1b* isoform. These results indicate that the proximal GLI-bound regions do not function, either individually or cooperatively, as major mesenchymal enhancers responsible for posterior *Ptch1* expression in the limb bud. In contrast, functional analysis of E335, E351, and E385 in transgenic reporter embryos revealed that, while E335 showed no detectable activity in limb buds, both E351 and E385 drove *lacZ* expression in the posterior limb mesenchyme (Fig. 5d; Fig. S8d). As E385 displayed robust posterior activity most closely resembling endogenous *Ptch1* expression, we next investigated if the function of this distal limb enhancer is evolutionarily conserved in artiodactyls. The orthologous pig and bovine E385 sequences retained enhancer activity in the posterior limb bud mesenchyme of transgenic reporter embryos (Fig. 5e). These analyses indicate that, while the LRM has undergone functional degeneration in both the pig and bovine lineages, the distal E385 enhancer retains limb enhancer activity in mouse reporter assays, which provides a likely explanation for the persistence of *Ptch1* upregulation, albeit restricted, in artiodactyl limb buds^1,3^ (Fig. 4).

### Cis-regulatory and pathway-level buffering preserve robust *Ptch1* upregulation in response to HH during limb development

The dispersed architecture of the *Ptch1* regulatory landscape predicts that limb expression is supported by enhancers with overlapping activities such as the LRM and the E351 and E385 elements. To test this hypothesis, we generated a series of targeted deletions affecting individual GLI-bound regions, alone or in combination (Fig. 6a). Thus, we engineered a deletion removing the GLI-bound element within the *Fancc* promoter (*Ptch1*^F^) and a large ∼300kb deletion (*Ptch1*^Δ300^) spanning the distal enhancer cluster, from E154 to E456, including the newly identified E351 and E385 limb enhancers (Fig. 6a; Fig. 5a). The E154 element was included in this large deletion because it displays weak regulatory activity in the limb^21^. These deletions were analyzed individually or combined in cis with the previously described *Ptch1*^AB^ allele (Fig. 4) to generate the Ptch1^F^, *Ptch1*^Δ300^, *Ptch1*^Δ300AB^ and *Ptch1*^Δ300ABF^ alleles (Fig. 6b). All alleles retained intact *Ptch1* promoter and coding sequences. Despite its chromatin interactions with the *Ptch1* locus (Fig. 5b), deletion of the GLI-bound region associated with the *Fancc* promoter did not produce detectable alterations in either the spatial distribution or expression levels of *Ptch1*, as assessed by WISH and bulk RNA-seq of posterior forelimb buds (Fig. 6b,c). Similarly, removal of the complete distal enhancer cluster in *Ptch1*^Δ300/Δ300^ embryos did not result in a statistically significant reduction of *Ptch1* expression at the posterior limb bud margin. In contrast, further weakening of the limb bud cis-regulatory repertoire in *Ptch1*^Δ300AB/Δ300AB^ embryos caused a significant decrease in *Ptch1* transcription, evident by both WISH and RNA-seq. The additional removal of the *Fancc*-associated GLI-bound region in *Ptch1*^Δ300ABF/Δ300ABF^ limb buds did not produce a statistically significant further reduction in *Ptch1* expression (Fig. 6b,c). None of these mutant embryos showed any evident forelimb skeletal patterning defects, and most of the additional HH pathway components showed no significant expression changes (Fig. S9a,c). Together, these experiments reveal an extensive functional redundancy across the *Ptch1* regulatory landscape, with distributed regulatory inputs buffering the effects of cis-regulatory deletions. The remarkable resilience of *Ptch1* transcription to extensive enhancer perturbation prompted us to investigate whether this robustness extends to the level of pathway regulation. PTCH1 occupies a unique position within the Hedgehog pathway, as it functions both as the receptor for the ligand and as a transcriptional target of pathway activation. Consequently, reduced PTCH1 dosage attenuates ligand sequestration, resulting in increased Hedgehog signaling that, in turn, stimulates *Ptch1* transcription through ligand-dependent feedback^13,15,19,20^. To experimentally assess this compensatory mechanism in embryonic limbs, we quantified *Ptch1* expression from the wild-type allele in *Ptch1*^ΔlacZ/+^ limb buds using an RT-qPCR amplicon that is absent in the knockout allele (Fig. 6d). Strikingly, expression from the remaining wild-type allele exceeded the 50% expected under a simple gene-dosage model and approached total *Ptch1* expression in wild-type limb buds (Fig. 6d). This finding is consistent with feedback regulation within the Hedgehog pathway buffering reductions in *Ptch1* dosage in limb buds through transcriptional compensation. Overall, our results indicate that the *Ptch1* locus is remarkably resilient to cis-regulatory perturbation, as reduced enhancer activity can be buffered both by regulatory redundancy and by pathway-level feedback, provided that sufficient Hedgehog-responsive regulatory regions remain to sustain transcriptional compensation.

**Figure 6.**
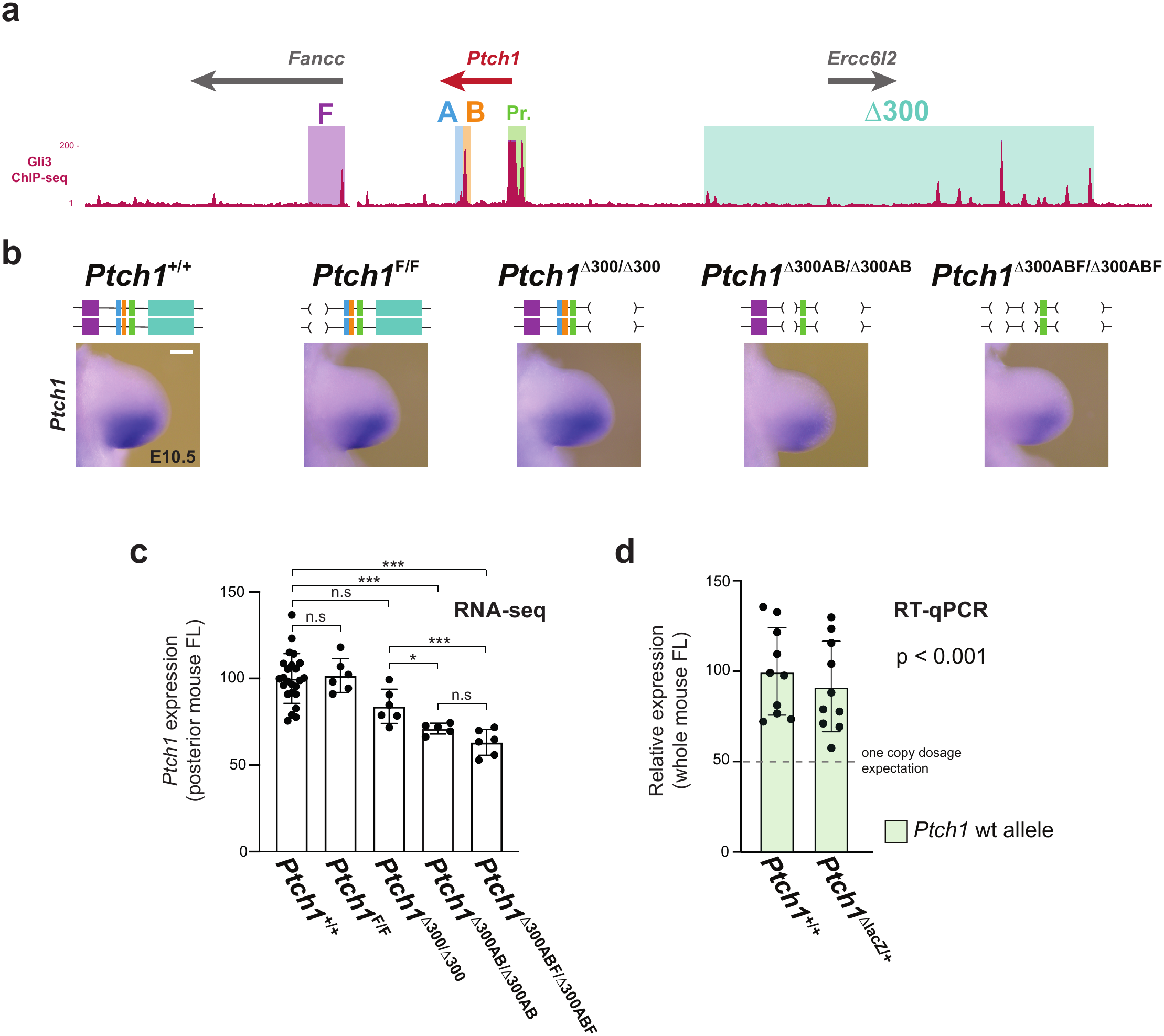
Functional dissection of the combinatorial cis-regulatory inputs that control *Ptch1* expression in mouse limb buds. (a) Schematic representation of the mouse *Ptch1* genomic landscape showing the neighboring genes *Fancc* and *Ercc6l2*, the promoter-proximal region (Pr.), and the deletions removing the most prominent GLI-binding regions, namely F (purple: ∼29 Kb deletion including the GLI-bound *Fancc* promoter), AB (deletion of regions A and B in the LRM, shown in blue and orange respectively) and Δ300 (cyan: a ∼300 Kb deletion removing all distal cis-regulatory regions between E154 and E456; see Figure 5). The GLI3 ChIP-seq profile from mouse limb buds is shown below^41^. (b) Whole-mount ISH for *Ptch1* in E10.5 mouse forelimb buds from the indicated genotypes (n=3 per genotype). While *Ptch1*^F/F^ and *Ptch1*^Δ300/Δ300^ mutant limb buds express *Ptch1* at levels comparable to those of wild-type controls, combined deletions progressively reduce *Ptch1* expression in mutant limb buds. Scale bar: 0.25mm. (c) RNA-seq quantification of *Ptch1* expression in posterior mouse forelimbs from the genotypes shown in (b). While *Ptch1*^F/F^ mutants show no statistically significant difference in *Ptch1* expression relative to wild-type controls, the *Ptch1*^Δ300^ mutation appears to decrease expression levels relative to *Ptch1*^+/+^ controls, although this trend does not reach statistical significance. In contrast, combining the *Ptch1*^Δ300^ deletion with the AB deletions (*Ptch1*^Δ300AB^) significantly reduces *Ptch1* expression compared with wild-type limb buds. Addition of the F deletion to this background (*Ptch1*^Δ300ABF^ allele) does not further reduce *Ptch1* expression to a statistically significant extent. Biological replicates are shown as individual dots. Statistical significance relative to corresponding wild-type littermates was assessed for each mutant genotype, and selected additional pairwise comparisons are indicated. For visualization, DESeq2 size-factor-normalized counts were transformed to relative expression values by setting the mean wild-type expression level to 100%. Data are shown as mean ± SD (n=6 per group, except n = 5 for *Ptch1*^Δ300^ homozygotes; all wild-type data points are shown). ***p ≤ 0.001; *p ≤ 0.05. P values correspond to BH-adjusted P values from DESeq2 Wald tests comparing each mutant genotype with the wild-type controls from the corresponding breeding line. For visualization purposes, the wild-type bar represents the pooled wild-type controls from all lines. FL: forelimb. n.s: not significant. Forelimb skeletal preparations and expression analyses of neighbouring genes and additional HH pathway components in this allelic series are shown in Fig. S9. (d) RT-qPCR analysis of *Ptch1* expression in E11.0 forelimbs from *Ptch1*^+/+^ and *Ptch1*^ΔlacZ/+^ embryos using primers spanning exons 2-3 which detect only the wild-type allele. Values are expressed relative to mean total *Ptch1* expression in wild-type embryos, set to 100%; the dashed line indicates the 50% expected from a single allele under a simple gene-dosage model. Dots represent individual embryos and bars show mean ± SD (n = 10 per genotype). Expression from the remaining wild-type allele in *Ptch1*^ΔlacZ/+^ limb buds was 90.9% of total *Ptch1* expression in wild-type embryos (95% CI, 71.3-116.0%) and significantly exceeded the 50% expected under a simple gene-dosage model (two-sided Welch’s t-test on log₂-transformed expression values, p=6.8 × 10^−5^.

## Discussion

Understanding how morphology evolves is particularly challenging as it relies largely on comparisons between species that are not all genetic model systems in developmental biology. Here, we show that single-cell multiomics can help bridge this gap by resolving transcriptional and regulatory differences within homologous embryonic cell populations of different species. As expected, mouse and pig limb buds share a broadly conserved cellular and regulatory organization, but display discrete differences in several morphoregulatory systems, including HOX-, FGF- and HH-modulated programs. These findings support the well-stablished view by which morphological novelties in evolution emerge through multiple changes distributed across otherwise deeply-conserved developmental networks, rather than through alterations in one or a few genes or enhancers^6,49^. This framework is particularly relevant to mammalian limbs, in which changes in early patterning and subsequent growth can jointly shape digit number, identity and skeletal proportions^1–5^. By comparing matched cell states rather than whole tissues, the approach also reveals expression and regulatory differences that would have been likely obscured by bulk measurements. Our paired mouse-pig atlas therefore provides both a resource and a strategy for using changes in gene regulatory network topology as entry points to approach evolutionary mechanisms.

One prominent difference we identified was the reduced expression of *FGFR2* and the simultaneous attenuation of FGF-pathway activity in the distal autopod mesenchyme of the pig. Pharmacological inhibition of FGFR signalling in mouse limb bud cultures reduced *Dusp6* expression and, similar to the genetic inactivation of *Fgfr1/2* in the mouse^26^ expanded the *Grem1* domain into the distal subectodermal mesenchyme, thereby recapitulating a characteristic feature of the pig *GREM1*. While these experiments do not establish reduced *FGFR2* expression as the sole cause of the pig-specific pattern, they demonstrate that attenuated FGF signalling is sufficient to phenocopy this spatial alteration. These findings position *GREM1* as an evolutionarily plastic node within the interconnected FGF, BMP and HH signalling networks^10,26,37,50,51^. *GREM1* spatial distribution could therefore be modified either through changes acting directly on its regulatory landscape or through alterations in upstream signalling inputs to help reshape distal growth and patterning. This interpretation is consistent with the known cis-regulatory plasticity of the *Grem1* locus and with recent studies linking changes in GREM1-mediated BMP antagonism to the spatial reorganization of distal mesenchymal populations^10,24^.

Although we focused primarily on the distal limb, our expression and regulatory analyses also identified differences in proximal pig limb mesenchymal populations. For example, the attenuation of HOXB2-HOXB6 regulons may contribute to the distinctive proportions of the artiodactyl proximal limb skeleton. This coordinated downregulation of several *HOXB* genes in pig could reflect evolutionary changes in shared cis-regulatory inputs within the *HOXB* genomic landscape^52^, analogous to the long-range regulatory domains that coordinately regulate multiple *HoxA* and *HoxD* genes in mouse limbs^53–55^. Although inactivation of *HoxB* genes has not been associated with major limb phenotypes in mice^56,57^, the conservation of their limb expression patterns across tetrapods suggests that these programs have retained functions that are not revealed by gross phenotypic characterization in mutant mice. In fact, inactivation of all *Hox5* paralogues in mice leads to severe proximal forelimb defects^58^. *Hoxb2-b6* genes may therefore act as secondary modifiers of limb growth or regional identity whose quantitative effects are difficult to detect, as previously shown for mice carrying mutations in *Fgfr2* or *Panx3* leading to subtle skeletal abnormalities revealed only through quantitative morphometric analysis^59,60^. Likewise, the functional significance of redeploying an otherwise conserved hindlimb-restricted *HOXC11* program in the pig distal forelimb remains uncertain. Severe zeugopod defects in mice develop after combined loss of *Hox11* paralogues, underscoring the paralogue redundancy that complicates inference from single-gene perturbations^61^.

These comparative analyses provide a starting point for examining how regulatory divergence can be tolerated at essential developmental loci. We therefore focused functional testing on the distal mesenchyme, where pig limb buds showed a broad shift towards posterior-like transcriptional states, consistent with previous evidence of altered anteroposterior polarity and changes in HH-responsive programs. Given its dual role as a direct SHH pathway target and negative-feedback regulator^13,15,19,20^, *Ptch1* provided a particularly tractable locus for testing genetically in the mouse how HH pathway output is encoded and evolutionary buffered against cis-regulatory change. Our results show that the functional degeneration of the ancient *Ptch1* LRM limb enhancer, previously identified in cattle, is shared with pigs. Whether LRM activity was lost independently after the divergence of these two early-branching artiodactyl lineages^62^ or was already absent in their last common ancestor remains unresolved. Nevertheless, its degeneration in both species suggests that partial erosion of *Ptch1* cis-regulatory robustness in the limb may be a defining feature in the evolution of the paraxonic artiodactyl autopod. Consistent with a contribution of the LRM to *Ptch1* regulation, *Ptch1*^Δ130/Δ130^ limb buds showed reduced *Ptch1* expression in the posterior limb bud mesenchyme, although the full effect cannot be attributed specifically to LRM ablation, as this deletion also removes additional minor GLI-bound regions in the interval and alters the native *Ptch1* transcript architecture. Next, our dissection of the broader *Ptch1* genomic landscape in mouse identified additional candidate limb enhancers distributed across the locus, the most prominent of which was E385, whose orthologous pig and bovine sequences retained limb enhancer activity in reporter assays in transgenic embryos. E385, together with additional untested sequences, could therefore sustain remaining *Ptch1* upregulation in the absence of the LRM, as in *Ptch1*^Δ130/Δ130^ embryos, or following its functional degeneration in artiodactyls.

The further genetic dissection of the *Ptch1* cis-regulatory landscape in the mouse revealed extensive functional redundancy, as individual deletions had little effect, whereas combining distal *Ptch1*^Δ300^ deletion with weakening of the LRM significantly reduced *Ptch1* transcription in the posterior limb margin. The remaining transcription may derive from residual LRM sequences or from other regulatory regions not removed by these deletions. We favor the former hypothesis because the LRM^ΔAB^ reporter retained residual limb mesenchymal enhancer activity, whereas the extended promoter E1:E8 reporter construct showed none; however, contributions from other untested regions cannot be excluded. This regulatory architecture resembles that of other pleiotropic developmental loci, in which partially redundant enhancers cooperate to maintain transcription above a functional threshold and thereby confer cis-regulatory robustness^7,9–11,53,63,64^. Such buffering could allow individual regulatory inputs to diverge while limiting the consequences for essential gene expression. Our allele-specific analysis revealed a second, pathway-level layer of buffering. Reducing functional *Ptch1* dosage increased transcription from the remaining wild-type allele, consistent with HH-dependent transcriptional compensation. *Ptch1* expression may therefore be stabilized through two complementary mechanisms, overlapping enhancer activity and signalling-dependent transcriptional feedback.

This feedback creates an apparent evolutionary paradox, as a cis-regulatory change that lowers *Ptch1* expression would attenuate LDA and thereby increase HH pathway activity, which would in turn stimulate compensatory *Ptch1* transcription through the remaining GLI-responsive cis-regulatory regions. Degeneration of the LRM in artiodactyls may nevertheless weaken a particularly important spatial and/or temporal input^65,66^, while enhancers such as E385 retain sufficient HH-responsive activity to sustain feedback. For such a change to be developmentally tolerated, it would need to reduce feedback-induced *Ptch1* expression sufficiently to alter ligand distribution while preserving enough basal PTCH1 activity to maintain LIA and avoid the severe defects associated with constitutive HH pathway activation^40^. Partial attenuation of individual cis-regulatory inputs may allow achieving this balance, together with additional mechanisms that remain to be characterized. Among these, the additional posterior domain of *HHIP* expression in pig limb buds may represent a further modification of the LDA system. Because HHIP binds and sequesters SHH^20,39^, this expression domain could provide additional spatial control over ligand distribution in the context of reduced PTCH1-mediated sequestration. For example, it could function as a localized ligand sink at the posterior margin, thereby reshaping the domain over which SHH remains available.

*Ptch1* provides a tractable example of how an essential developmental locus can remain both robust and evolvable through the combined action of enhancer redundancy and signalling feedback, and similar nested layers of cis-regulatory and pathway-level buffering may operate in other morphoregulatory networks. More broadly, our findings illustrate how comparative gene regulatory network inference at cell-type resolution can reveal distributed regulatory divergence within deeply conserved developmental systems and guide functional analyses of how essential loci accommodate regulatory change.

## Methods

### Mouse and pig embryos

Mouse (*Mus musculus*) and domestic pig (*Sus scrofa domesticus*) embryos were collected in accordance with Spanish and EU regulations, and all animal experimentation was performed in strict compliance with the 3R principles. Wild-type CD1 mouse embryos at 10.5 or 11.5 days post coitum (E10.5-E11.5) were obtained by natural mating. Pig embryos at 21 or 23 days post coitum (D21-D23) were obtained after hormonal synchronization of recipient sows followed by natural mating. The mating date was considered gestational day 0 (D0), and pregnancy was confirmed by ultrasound. The overall production time for each litter was approximately 45-50 days, once the sows had reached a minimum age of 8-9 months. The equivalence between mouse and pig limb developmental stages was established in previous studies^3^.

### Whole-mount *in situ* hybridization of mouse and pig limb buds

Whole-mount in situ hybridization (WISH) was performed on dissected mouse (E10.5-E11.5) and pig (D21-D23) limb buds as previously described^3^. Embryos were collected in ice-cold PBS, fixed overnight in 4% PFA at 4 °C, dehydrated into methanol and stored at −20 °C. All embryos were split so that left and right limb buds could be processed separately. Briefly, samples were rehydrated, bleached in 6% H_2_O_2_ and permeabilized with proteinase K under stage-specific conditions (mouse E10.5, 15 min; mouse E11.5, 25 min; pig D21, 35 min; pig D23, 45 min). After postfixation, samples were prehybridized for 1 h and hybridized overnight at 70 °C with DIG-labelled riboprobes. Following high-stringency washes and RNase A treatment, samples were blocked in 10% lamb serum, incubated overnight at 4 °C with anti-DIG-AP antibody (1:5,000; Roche), and developed in BM Purple (Roche). Reactions were stopped by washing in PBT, and samples were post-fixed in 4% PFA/PBS. Images were acquired using a Leica M125 C stereomicroscope equipped with a Leica Flexacam C1. Brightness and contrast were adjusted linearly using Adobe Photoshop or ImageJ. For each probe, a minimum of three mouse or two pig limb buds were analyzed per stage. In selected cases, owing to the limited availability and high value of pig embryonic material, one replicate for the earlier stage (D21) corresponded to the hindlimb of a D23 embryo, as previously described, taking into account the developmental delay between forelimbs and hindlimbs^3^. The sequences of all mouse and pig WISH probes generated for this study are provided in Table S9 and were generated by PCR from limb bud cDNA or synthesized as DNA fragments, and cloned into pBluescript KS(+). Mouse and pig probes were designed to target the same conserved exonic regions. All pig riboprobes were first validated in mouse embryos.

### Differential expression analysis by RT-qPCR and bulk RNAseq

Forelimb buds were dissected in ice-cold PBS and stabilized in RNAlater before storage at −20°C. Total RNA was extracted from samples of the appropriate genotype, reverse-transcribed into cDNA, and analyzed by RT-qPCR as described previously^67^. Relative transcript levels were normalized to *Rpl19* and *Hprt1*, and expression in mutant samples was calculated relative to wild-type controls whose mean was set to 100%. Data are shown as mean ± SD from 8-10 biological samples (independent embryos) per genotype. Statistical significance was assessed using an unpaired *t*-test. The hypothesis that *Ptch1* expression in heterozygous limb buds from the remaining wild-type allele exceeded the expected 50% was tested on log_2_-transformed values using a two-sided unpaired Welch’s t-test. Statistical analysis was performed with Prism 9. The sequences of the qPCR primers used are provided in Table S10.

For RNA-seq, the posterior halves of left and right E11.0 forelimb buds were dissected, pooled and preserved in RNAlater at −20°C. After genotyping and sex determination, six samples per genotype (three males and three females) were selected for total RNA extraction and sequencing, together with six somite- and background-matched wild-type controls. Poly(A)+ RNA was used for library preparation with the Optimal Dual-mode mRNA Library Prep Kit (BGI), and libraries were PE100 sequenced on the DNBSEQ-G400 platform (BGI Genomics). Initial processing of RNA-seq data was performed by BGI. Raw reads containing adaptor sequences or low-quality bases were filtered using SOAPnuke with the parameters -n 0.01 -l 20 -q 0.4 --adaMR 0.25 --polyX 50 --minReadLen 100. Reads were discarded if they matched adaptor sequence over ≥25% of their length, were shorter than 100 bp, contained >1% ambiguous bases, contained polyX stretches longer than 50 bp, or had ≥40% bases with quality score <20. Clean reads were provided in Phred+33 format.

Clean paired-end reads were quality-controlled using FastQC v0.12.1 and aligned to the mouse GRCm38/mm10 reference genome using HISAT2 v2.2.1 with reverse-forward strand-specific settings (--rna-strandness RF). SAM files were converted to sorted BAM files and indexed using SAMtools v1.21. Gene-level counts were generated with featureCounts v2.0.8 from the Subread package and the GENCODE vM10 mouse gene annotation (gencode.vM10.annotation.gtf). featureCounts was run with the options -p --countReadPairs - s 2 -B -C, corresponding to paired-end fragment counting, reverse-stranded libraries, requiring both mates to be successfully aligned, and excluding chimeric fragments. FastQC and featureCounts outputs were summarized using MultiQC v1.28.

Individual gene-level count files were merged by Ensembl gene identifier to generate a gene-by-sample count matrix. One sample was excluded before the final differential expression analysis because of misgenotyping, resulting in a final dataset of 47 samples. Differential expression analysis was performed in R v4.4.1 using DESeq2 v1.44.0^68^. Raw counts were imported with sample metadata, and a DESeq2 dataset was constructed using genotype as the design factor (∼ Genotype). Wild-type samples (*Ptch1*^+/+^) were set as the reference level. Genes with fewer than 10 total counts across all samples were removed, retaining 23,889 genes for analysis. Size-factor normalization, dispersion estimation and negative binomial generalized linear modelling were performed using the standard DESeq2 workflow. *Ptch1* differential expression was tested for *Ptch1*^F/F^, *Ptch1*^Δ300/Δ300^*, Ptch1*^Δ300AB/Δ300AB^ and *Ptch1*^Δ300ABF/Δ300ABF^ mutant samples relative to their corresponding wild-type controls using Wald tests. Pairwise comparisons between mutant genotypes were also performed by generating separate DESeq2 datasets containing only the two genotypes being compared. P values were adjusted using the Benjamini-Hochberg (BH) procedure, and genes with adjusted P < 0.05 were considered differentially expressed. As a sensitivity analysis, the same comparisons were repeated after combining all wild-type controls into a single reference group, yielding similar results.

### Enhancer reporter assays in transgenic mouse embryos

Transient transgenic mouse embryos were generated by pronuclear injection at CTM-University of Basel (mouse LRM^ΔAB^ and mouse E335, E351 and E385), the LBNL (pig LRM, pig E385, bovine E385 and mouse E3) or CABD (E1:E8). Coordinates of the assayed regions are indicated in Table S11. Injected embryos were transferred into 0.5 dpc pseudopregnant females, collected at E10.5-E11.75, and genotyped by PCR using *lacZ* primers. Embryos injected with the mouse, pig and bovine LRM constructs at CTM-Unibasel were carried to term to establish stable transgenic lines, with their expression patterns matching those observed in the transient transgenic embryos. For β-galactosidase staining, embryos were fixed for 20 min in fixative solution (1% formaldehyde, 0.2% glutaraldehyde, 0.04% IGEPAL CA-630 and 0.01% sodium deoxycholate in PBS), rinsed three times for 5 min in PBS, and incubated in staining solution (0.5 mg/ml X-gal, 250 nM K_3_Fe(CN)_6_, 250 nM K_4_Fe(CN)_6_, 400 nM MgCl_2_, 0.01% IGEPAL CA-630 and 0.01% sodium deoxycholate in PBS). The reaction was stopped by extensive washing in PBS, followed by postfixation in 4% PFA in PBS. E4, E11 and E60 reporter transgene expression was previously reported in the VISTA database^45,46^.

### Limb-bud culture and pharmacological inhibition

Mouse embryonic forelimbs from wild-type C57BL/6J E10.5 embryos were dissected with sufficient trunk tissue to allow pinning and cultured *in vitro* as previously described^36,51^. Explants were maintained for 16h at 37 °C in 6% CO2 in serum-free high-glucose DMEM supplemented with penicillin/streptomycin, L-glutamine, non-essential amino acids, sodium pyruvate, D-glucose, L-ascorbic acid, lactic acid, d-biotin, vitamin B12 and PABA. FGF signalling was inhibited with SU5402 (10 μM final concentration; Stem Cell Technologies), and control cultures supplemented with 0.1% DMSO as vehicle. At the end of the culture period, samples were rinsed in PBS and fixed overnight in 4% PFA/PBS at 4 °C. Experiments were repeated with at least three independent embryos, with equivalent results.

### UMI-4C-seq

For each replicate, forelimb buds of the same developmental stage (E11.0; 40-42 somites) were dissected in cold PBS and pooled to obtain sufficient material per sample (30-35 embryos per replicate). Tissue was washed in 10% FBS/PBS and enzymatically dissociated with collagenase at a final concentration of 0.8 mg/ml in prewarmed 10% FBS/PBS. Limb buds were first incubated for 10 min at 37 °C with shaking at 750 rpm, after which the supernatant was passed through a prewetted 70 µm cell strainer. The remaining tissue was subjected to a second collagenase digestion at 37 °C with intermittent vigorous trituration until complete dissociation, and the resulting cell suspension was filtered through the same 70 µm strainer. Filters were washed with 10% FBS/PBS and cells counted. Cell suspensions (8.7-9.6 × 10^6^ cells) were adjusted to a final volume of 11.4 ml with 10% FBS/PBS and crosslinked by addition of methanol-free formaldehyde to a final concentration of 2% for 10 min at room temperature with gentle rocking. Crosslinking was quenched with glycine to a final concentration of 0.13 M, and samples were transferred to ice. Cells were pelleted by centrifugation at 400g for 8 min at 4 °C, washed with cold PBS, and lysed in cold lysis buffer (50 mM Tris-HCl pH 7.5, 150 mM NaCl, 5 mM EDTA, 0.5% IGEPAL CA-630, 1% Triton X-100 and 1x cOmplete protease inhibitor cocktail) for 10 min on ice. Nuclei were collected by centrifugation at 500g for 5 min at 4 °C, transferred to safe-lock tubes, briefly pelleted at 500g, snap-frozen in liquid nitrogen and stored at −80 °C until library preparation. Nuclei preparations were digested with DpnII and proximity-ligated *in situ* using T4 DNA ligase. Crosslinks were reversed by overnight incubation with Proteinase K, followed by RNase A treatment and DNA purification to generate the 3C template. UMI-4C libraries were prepared as described^47^. Briefly, 3C templates were sonicated to an average fragment size of approximately 500 bp, end-repaired, A-tailed and ligated to Illumina-compatible forked adapters containing unique molecular identifiers (UMIs). Viewpoints were amplified by two rounds of nested PCR using locus-specific upstream and downstream primers listed in Table S12. UMI-4C amplicons were purified using AMPure XP beads, quantified and pooled for paired-end (PE50) sequencing in a DNBSEQ-G400 system (BGI Genomics). Data analysis was performed in R v4.4.1 using UMI4Cats v1.14.0^69^. An in silico DpnII-digested mouse reference genome was first generated from the mm10 genome using the digestGenome() function, with the DpnII recognition sequence GATC and cut position 0. Multiplexed FASTQ files were demultiplexed by viewpoint using the corresponding bait/barcode sequences, and viewpoint-specific FASTQ files were organized for mapping. For each viewpoint, UMI-supported contacts were generated with the contactsUMI4C() function using the viewpoint-specific bait and padding sequences, the in silico-digested mm10 genome and a Bowtie2 mm10 index. Processing statistics were summarized with the statsUMI4C() function. Per-viewpoint UMI4C objects were generated from the contact-count tables using the makeUMI4C() function, with bait_expansion = 3e6, min_win_factor = 0.01.

### Generation and analysis of engineered mouse alleles

The *Ptch1*^ΔlacZ^ and Shh^Δ^ alleles have been described previously^43,70^. The *Ptch1*^A^ and *Ptch1*^B^ alleles, which delete the two GLI-binding regions within the *Ptch1* LRM, were generated individually in G4 mouse embryonic stem (mES) cells^71^ by CRISPR/Cas9-mediated genome editing, as described previously^72^ using the px459 plasmid^73^. The sgRNA sequences used to induce these deletions, together with the primers used for genotyping, are listed in Table S13. Correctly targeted ES cell clones were identified by PCR and confirmed by sequencing, and mutant mouse lines were established by aggregation chimeras with diploid embryos. The double-mutant *Ptch1*^AB^ allele was generated by re-targeting ES cells carrying the A deletion and proven germline transmission potential with sgRNAs directed against the B region, followed by screening for deletions occurring in *cis* on the same chromosome. The *Ptch1*^Δ130^ allele was generated in G4 ES cells by co-transfection of px459 plasmids encoding sgRNAs flanking the targeted region (Table S13) together with a repair template harbouring an artificial *Ptch1* exon 13-23/BGHpA cassette and the endogenous intronic sequences containing the corresponding donor splice site, all flanked by ∼700-bp homology arms. Homozygous *Ptch1*^Δ130/Δ130^ ES cells were used to generate aggregation chimeras with tetraploid embryos, thereby ensuring that the embryo proper was fully derived from the targeted ES cells^44^. Control embryos were obtained from wild-type matings of the same genetic background. All other alleles generated in this study were produced by zygote electroporation of in vitro assembled Cas9/tracrRNA/crRNA RNPs^74^ (Table S13). The *Ptch1*^ΔPrP^, *Ptch1*^F^ and *Ptch1*^Δ300^ alleles were generated on a CD1 wild-type background, while the *Ptch1*^Δ300AB^ and *Ptch1*^Δ300ABF^ alleles were obtained by electroporation of zygotes carrying the *Ptch1*^AB^ or *Ptch1*^Δ300AB^ alleles in a CD1 background with RNPs containing Δ300 or F crRNAs, respectively. The resulting mice were screened for in-*cis* targeting events, and the targeted rearrangements were confirmed by sequencing. All mutant mouse lines were maintained on a CD1 background (Janvier, France) for >10 generations by serial outcrossing of heterozygotes to wild-type CD1 animals obtained from the same supplier. Experimental cohorts were generated by heterozygous intercross and analyzed against somite-matched littermate controls. Embryos of both sexes were used in the analyses.

### Nuclei isolation from mouse and pig limb buds for single-cell multiome assays

Single-nucleus suspensions for single-cell Multiome assays were prepared following an adapted version of the 10x Genomics demonstrated protocol Nuclei Isolation from Embryonic Mouse Brain for Single Cell Multiome ATAC + Gene Expression Sequencing (CG000365, Rev. C). Unless otherwise indicated, all steps were performed on ice and centrifugations were carried out at 300g for 5 min at 4 °C. Centrifuge rotors and dissection tools were thoroughly cleaned with sterile Milli-Q H2O and treated in advance with RNaseZap (Ambion). For all filtration steps, cell strainers were pre-wetted with a small volume of the corresponding sample-containing buffer before use.

Mouse (CD1 strain) and pig embryos were collected in ice-cold PBS for staging (mouse, E10.5 and E11.5; pig, D21 and D23), and forelimb buds were dissected together with the adjacent flank mesenchyme in order to capture proximal progenitors contributing to the shoulder girdle. Five E10.5 mouse embryos (35-36 somites; 10 forelimb buds), two E11.5 mouse embryos (45-46 somites; 4 forelimb buds), four D21 pig embryos (8 forelimb buds) and two D23 pig embryos (4 forelimb buds) were processed. Forelimb buds were pooled within each species and developmental stage, yielding four stage-specific multiome samples. Individual embryo identities were not retained after pooling. Freshly dissected forelimbs were washed three times in D-PBS (Gibco), once in 0.4% bovine serum albumin (BSA; ThermoFisher, AM2616) in D-PBS, and transferred to a 2-ml dounce tissue grinder (Kimble, 885300-0002). Tissue was mechanically dissociated into a near single-cell suspension using the 127-76 μm pestle (pestle A, 885301-0002). Dissociation was achieved with six strokes for mouse limb buds and ten strokes for pig limb buds. These conditions were adjusted for pig samples, as pig limb tissue is notably more resistant to mechanical dissociation. The resulting suspension was recovered into a 1.5ml Eppendorf tube and, to maximize recovery, the homogenizer was rinsed with 0.4% BSA in D-PBS and the rinsate was combined with the original suspension. After centrifugation (300g, 4 min, 4 °C), the supernatant was carefully aspirated and the pellet resuspended in 500μl lysis buffer containing 10 mM Tris-HCl, pH 7.4, 10 mM NaCl, 3 mM MgCl2, 1% BSA (Thermo Fisher), 0.1% Tween-20 (Bio-Rad), 0.1% IGEPAL (Sigma), 0.01% digitonin (Thermo Fisher Scientific), 1 mM DTT and 1 U/μl RNase inhibitor (Roche). Efficient lysis was achieved by gentle pipetting with a P1000 tip for 1 min (mouse samples) or 1 min 30 s (pig samples). Pipetting for longer than 2 min was avoided in order to preserve nuclear membrane integrity. Lysis was quenched by addition of 900μl wash buffer containing 10mM Tris-HCl, pH 7.4, 10mM NaCl, 3mM MgCl2, 1% BSA (Thermo Fisher), 0.1% Tween-20 (Bio-Rad), 1mM DTT and 1 U/μl RNase inhibitor (Roche). The lysate was passed through a 70-μm cell strainer, followed by an additional 1 ml wash buffer through the filter to maximize sample recovery. To minimize nuclei aggregation, the suspension was divided equally into two 1.5ml Eppendorf tubes before the next centrifugation step. The resulting nuclei pellets were each resuspended in 400 μl wash buffer and then filtered through a 45-μm strainer, with the flow-through collected into a single tube. After centrifugation, the final pellet was resuspended in 200-300 μl diluted nuclei buffer, with the volume adjusted according to the amount of starting material (1× Nuclei Buffer, 10x Genomics, supplemented with 1 mM DTT and 1 U/μl RNase inhibitor, Roche). Nuclei concentration was determined using a Neubauer chamber, and nuclei integrity was assessed by DAPI staining followed by fluorescence microscopy. The percentage of live cells was below 3% in all samples (TC20 cell counter; Bio-Rad).

### Single-cell multiome library preparation and sequencing

Single-cell multiomic (ATAC + gene expression) libraries were prepared by the Functional Genomics Facility at the CABD using the 10x Genomics Chromium Controller and the Chromium Next GEM Single Cell Multiome ATAC + Gene Expression Reagent Bundle (PN-1000285). For each sample, ∼20,000 nuclei were subjected to transposition and loaded onto a Chromium Next GEM Chip J (Single Cell Kit, PN-1000230) for partitioning of single nuclei into gel beads-in-emulsion. Barcoding and pre-amplification were performed using the Single Index Kit N Set A (PN-1000212) and the Dual Index Kit TT Set A (PN-1000215), according to the manufacturer’s instructions (CG000338 Rev. C). cDNA and ATAC library concentrations were measured using the Qubit dsDNA High Sensitivity Assay on a Qubit 4 Fluorometer (Invitrogen), and library size profiles were assessed using a Fragment Analyzer before sequencing. Libraries were sequenced on a DNBSEQ-G400 system (BGI Genomics) using PE100/100/10/10 settings for cDNA libraries and PE50/50/8/16 settings for ATAC libraries.

### Computational analyses of scRNA- and scATAC-seq (multi-omics) data

#### Pre-processing

Reads were aligned to the GRCm38/mm10 mouse or Sus scrofa 11.1 pig genome assemblies. Cell Ranger ARC (v.1.0.0). recovered 7,975 cells at E10.5, with a median of 3,126 genes per cell (cDNA, 39,617 mean raw reads per cell; ATAC, 72,393 mean raw reads per cell); 6,770 cells at E11.5, with a median of 2,370 genes per cell (cDNA: 39,917 mean raw reads per cell; ATAC: 60,271 mean raw reads per cell); 17,499 cells at D21, with a median of 2,267 genes per cell (cDNA: 30,031 mean raw reads per cell; ATAC: 45,610 mean raw reads per cell); and 5,321 cells at D23, with a median of 3,001 genes per cell (cDNA: 51,910 mean raw reads per cell; ATAC: 77,519 mean raw reads per cell).

Pre-processing was performed with Seurat (v.4.0.3)^75^ and Signac (v.1.2.1)^48^ R packages. As a quality control, for both species we filtered out the cells that fulfilled any of the following conditions: <200 RNA counts; >40,000 RNA reads; >7,000 detected genes; > 100,000 ATAC reads; <500 ATAC reads; TSS enrichment score < 2; nucleosomal signal score >2.5. For the mouse we filtered cells with >10% mitochondrial RNA reads and <800 detected genes, while for the pig we filtered cells with >5% mitochondrial RNA reads and <700 detected genes. For each species, we merged the different scATACseq libraries creating a common set of peaks, with the UnifyPeaks() Signac function. Transcript and peak counts were normalized using log and tf-idf normalization, respectively. To remove unwanted variability, we filtered out a set of housekeeping, cell cycle, and ribosomal genes from the RNA count matrix.

The pre-processing of the samples consisted of the following steps: we identified the highly variable genes (HVG) using the FindVariableFeatures, followed by data scaling, and Principal Component Analysis (PCA) using the default parameters of ScaleData() and RunPCA() functions of Seurat. The different libraries of each species were integrated using the Harmony software (v.0.1.0)^76^, following a workflow previously applied to single-cell datasets spanning developmental stages of mouse endochondral bone^77^, using the first 30 principal components of the RNA processed data.

#### Clustering and cell annotation

We clustered the cells based on 30 Harmony RNA dimensions, using the FindNeighbors() and FindClusters() Seurat functions. The non-mesenchymal cells separated clearly from the mesenchymal cells and were annotated based on canonical gene markers. For a careful annotation of mesenchymal cell types we used the mouse dataset as reference. We first subset the mouse mesenchymal cells from the rest of the cells for pre-processing and clustering following the same approach mentioned above. The mouse mesenchymal clusters were annotated based on the markers shown in Figure S1, resulting in six mesenchymal clusters (“DIM”, “PIM”, “PFM”, “DAM”, “PPM”, “GHM”). We then annotated the homologous pig mesenchymal cell types using label transfer (projection of reference metadata onto the query object). For this we subset the expression matrix of both species to include only 1-1 ortholog genes. Then we used the Seurat functions FindTransferAnchors() and MapQuery() setting PCA as the reference reduction. Finally, to annotate the Shh cluster (“ZPA”) in each species, we selected cells from the distal mesenchyme (clusters “DAM” and “DIM”) with an expression level of at least 2 counts for the *Shh*/*SHH* gene.

#### Peak calling and re-quantification of chromatin accessibility

To refine chromatin accessibility profiles, we re-called peaks from the scATAC-seq data using the CallPeaks() function in Signac, which implements the MACS3 algorithm (v3.0.0a6)^78^. Peak calling was performed independently for the mesenchymal datasets from mouse and pig, and within each species, separately for every annotated mesenchymal cluster (group.by = “Anno_180324”) to capture cluster-specific open chromatin regions. As implemented in Signac, CallPeaks merges all identified peaks across clusters into a single, non-redundant set of genomic intervals, retaining all unique peaks within the dataset. The resulting peak sets were filtered to remove non-standard chromosomes using keepStandardChromosomes(pruning.mode = “coarse”). Peaks overlapping ENCODE blacklisted regions were excluded in the mouse dataset (subsetByOverlaps, invert = TRUE), a step not performed for pig due to the lack of a validated blacklist^79^. Finally, fragment counts overlapping the MACS3-derived peaks were re-quantified using FeatureMatrix. This redefined “peaks” assay was subsequently used for genomic feature annotation (RegionStats), peak-to-gene correlation analysis (LinkPeaks) and peak-to-gene visualization (CoveragePlot). The peak-to-gene links derived from this analysis were also exported to the UCSC Genome Browser by generating *.interact* files.

#### Identification of evolutionarily divergent limb mesenchymal genes

We performed cell-level differential expression analyses between homologous mesenchymal clusters in mouse and pig, as defined by the label transfer procedure described above. Cells from the two stage-specific pooled libraries for each species were analyzed together. As embryos identity was not retained in library preparation, differential expression was assessed at the cell level. Adjusted P values therefore quantify cell-level differences and do not estimate between-embryo variability. These analyses were used to identify candidate interspecies expression differences, selected examples of which were assessed spatially by WISH. To ensure cross-species comparability, we followed these steps: (i) pig FASTQ files were remapped to a reference in which the genomic coordinates for *HOXD9*, *HOXD11*, and *HOXD12* were manually curated and extended, as the previous reference led to incomplete assignment of reads from these loci to the annotated gene bodies; (ii) we retained the previously computed dimensionality reduction and clustering, as described above; and (iii) NormalizeData() was re-run after restricting the mouse and pig datasets to the same 1:1 ortholog set, so that per-cell library sizes (total counts) used for normalization were computed over a matched gene space and were not confounded by species-specific differences in GTF gene models. We then used Seurat (v 5.4.0) FindAllMarkers() function (min.pct = 0.1; |log_2_ fold change| ≥ 1) to identify inter-species differentially expressed genes (DEGs) for each homologous cluster (e.g., mouse ZPA vs. pig ZPA), retaining only genes with adjusted *P* value < 0.05. These cell-level comparisons were used to prioritize candidate expression differences between the sampled populations. To enrich for cluster-associated differences rather than broadly distributed species differences, we further filtered the set of DEGs for each cluster to retain only those genes that were also identified as within-species cluster markers in at least one of the two species (e.g., mouse ZPA vs. mouse non-ZPA), under the same statistical thresholds. Genes located on sex chromosomes and genes lacking an associated Entrez ID were excluded from all downstream analyses. Next, we used Gephi to generate graphs in which each mesenchymal cluster is connected to its corresponding DEGs, visualizing only genes associated with limb phenotypes and using node color to indicate whether each gene is upregulated in mouse (blue) or in pig (red). The list of limb phenotype (LP) genes (Table S4) was generated by merging the protein coding genes included in the GO:0060173 (limb development), MP:0002109 (abnormal limb morphology) and MP:0009250 (abnormal appendicular skeleton morphology) categories, and manually curated to add the *Meis1* and *Meis2* genes, with essential functions in limb development^80^.

#### Functional enrichment analyses of evolutionarily divergent limb mesenchymal genes

To characterize the biological processes associated with evolutionary divergence, we performed functional enrichment of Gene Ontology^81^ (GO) terms on the evolutionarily divergent limb mesenchymal gene set for each cluster. Analyses were carried out using the Bioconductor packages clusterProfiler^82^ (v.4.18.4), together with the mouse-specific annotation database org.Mm.eg.db (v. 3.22.0; https://doi.org/10.18129/B9.bioc.org.Mm.eg.db). We used the function enrichGO() with the default parameters (pAdjustMethod = “BH”, pvalueCutoff = 0.05, and qvalueCutoff = 0.2), all of which implement a hypergeometric test^83^.

#### DAM subclustering into anterior and posterior states

For each species, DAM cells were subsetted into a separate Seurat object and reprocessed independently. All genes, rather than only orthologues, were included in this analysis. Harmony batch correction was rerun using the PCA space inherited from the full mesenchymal object, rather than from the DAM-only subset, as this representation best resolved the anterior-posterior axis. The Harmony embedding was then used for UMAP visualization, nearest-neighbor graph construction, and low-resolution clustering, yielding two DAM subclusters. Based on canonical marker expression, these subclusters were annotated as anterior and posterior. The refined DAM labels were then projected onto the original mesenchymal UMAP.

#### SCENIC+

We applied SCENIC+^27^ independently to mouse and pig datasets to infer enhancer-driven gene regulatory networks (eGRNs) by integrating scRNA-seq, scATAC-seq, and motif discovery. The workflow comprises three modules: pycisTopic, pycisTarget, and the SCENIC+ Snakemake pipeline:

##### pycisTopic

We used pycisTopic (v.2.0a0) to analyze the scATAC-seq data and identify candidate enhancers. This comprised: (i) generation of pseudobulk profiles from cell clusters; (ii) peak calling with MACS3 for each pseudobulk profile; (iii) derivation of consensus peaks using an iterative filtering approach^84^ (*peak_half_width* = 250); (iv) computation of QC metrics at both sample and barcode levels; and (v) construction of a count matrix of fragment counts over consensus peaks for all barcodes passing QC. For mouse data, regions overlapping the mm10 blacklist (mm10-blacklist.v2.bed) were removed, a step omitted for pig due to the lack of a validated *Sus scrofa* blacklist^79^. Topic modeling was performed using parallel latent Dirichlet allocation implemented in MALLET (https://mimno.github.io/Mallet/index). The number of topics (45 for each species) was selected following pycisTopic guidelines, prioritizing topic coherence^85^ and final-iteration log-likelihood^86^. Putative regulatory regions were defined by combining binarized topic-region distributions (via Otsu and ntop methods) and cell cluster-specific differentially accessible regions (DARs).

##### pycisTarget

pycisTarget (v.1.0a2), run within the SCENIC+ Snakemake pipeline, was used for motif enrichment to identify potential transcription factor binding sites (TFBS) in candidate enhancers, implementing the cisTarget ranking-and-recovery algorithm and the DEM (differential enrichment of motifs) Wilcoxon rank-sum test. For each species, we provided a cisTarget database built by scoring motifs across pycisTopic consensus regions with cbust^87^, using the v10nr_clust_public motif collection (https://resources.aertslab.org/cistarget/motif_collections/v10nr_clust_public/) and a motif-to-transcription factor (TF) annotation table (SCENIC+ mouse resource for mouse; custom orthology-derived annotation for pig).

##### SCENIC+ Snakemake pipeline

We used the SCENIC+ Snakemake pipeline to integrate the scRNA-seq data with the previous outcomes and reconstruct eGRNs. The scRNA-seq count matrix was preprocessed according to SCENIC+ guidelines, including normalization and scaling. Some configuration parameters in the *config.yaml* file were customized. Specifically: (i) the extended motif annotation was based on orthology and/or motif similarity, thus enabling broader regulatory network reconstruction in species with comparatively sparse regulatory annotations, such as the pig (default: orthology only); (ii) the maximum search space around each gene was extended to 300 kb upstream and downstream of the gene body (default: 150 kb); (iii) for each gene, regulatory regions and transcription factors were independently ranked based on rho metrics, rather than importance (default); and (iv) the minimum number of target genes required to define an eRegulon was set to 8 (default: 10).

As part of the SCENIC+ pipeline output, we obtained TF-region-gene triplets organized into eRegulons, each defined by a transcription factor and its associated set of regulatory regions and target genes (Tables S7 and S8). A single TF can give rise to multiple eRegulons, which are characterized by the sign (positive or negative) of two key correlation metrics: R2G_rho, reflecting the relationship between chromatin accessibility at the region and expression of the target gene, and TF2G_rho, capturing the co-expression between the TF and its target gene. In addition, we obtained eRegulon activity at single-cell resolution (computed with AUCell), as well as eRegulon specificity scores (RSS) for each cell cluster. In our downstream analysis, eRegulons with negative R2G_rho were excluded, as well as those involving FREM1 as a TF, as it was erroneously included in the SCENIC+ transcription factor list for mouse (https://resources.aertslab.org/cistarget/tf_lists/allTFs_mm.txt). To construct cluster-specific regulatory networks with Gephi^88^ (v 0.10.1), we selected, for each cluster, the eRegulons with the highest RSS scores until reaching a total of 20 distinct TFs. Only activator eRegulons (those with positive TF2G_rho) were considered in this selection since eGRN inference is still biased toward activation and remains less reliable for identifying repressive interactions^27^. Both the regulatory networks and the *coverage plots* visualizations were generated using triplets with R2G_rho ≥ 0.1 (default threshold: 0.05). The regulatory links shown in the coverage plots were exported to the UCSC Genome Browser via the creation of *.interact* files.

## Use of artificial intelligence tools

During preparation of this manuscript and data analysis, ChatGPT was used to assist with language editing, including corrections to grammar and style, and with drafting, reviewing and debugging analysis code. All AI-edited code was reviewed, adapted and executed by the authors, who verified its operation and take full responsibility for the analyses and resulting outputs.

## Data availability

All sequencing data generated in this study will be deposited in the GEO and accession numbers will be provided prior to publication.

## Code availability

Scripts necessary to reproduce our analysis will be publicly available in a GitHub repository upon publication.

## Acknowledgements

We wish to thank A. Franco, C. Mateos, A. López and P. López for mouse husbandry and all personnel at the Servicio de Experimentación Animal (SAEX; Universidad de Córdoba) for pig care. We are also grateful to Rafael Rodríguez Acemel and Ildefonso Cases for bioinformatic advice.

## Funding

This work was supported by the Spanish Agencia Estatal de Investigación (MCIN/AEI/ 10.13039/501100011033 / FEDER, UE) through grants PID2020-113497GB-I00 and PID2023-148267NB-I00 to J.L.R, PID2022-141288NB-I00 to J.J.T., PID2022-143253NB-I00 to D.G.L., RYC2022-035182-I and PID2023-151163NA-I00 to F.M.R and PID2022-137086NB-I00 to B.P. D.G.L is also supported by the European Research Council (grant no. 101045439, 3D-REVOLUTION). Funded by the European Union. Views and opinions expressed are however those of the author(s) only and do not necessarily reflect those of the European Union or the European Research Council Executive Agency. Neither the European Union nor the granting authority can be held responsible for them. M.O was supported by Swiss National Science Foundation (SNSF) grants PCEFP3-186993 and 10008859. A.V. was supported by NIH grant R01HG003988. Research conducted at the E.O. Lawrence Berkeley National Laboratory and performed under U.S. Department of Energy Contract DE-AC02-05CH11231, University of California (UC).

## Author contributions

Conceptualization: G.N., J.L.R.; Methodology: F.B., G.N., J.L.R.; Software: I.S.M., L.A.; Formal Analysis: A.A.C., I.S.A., V.T., I.S.M., L.A., J.P., G.N., J.L.R.; Investigation: A.A.C., I.S.A., V.T., A.D.R., F.B., A.F.M., B.J.M., A.H.M., F.M.R., B.P., M.O., G.N., J.L.R.; Resources: D.G.L., B.P., M.O., A.V., H.H., J.J.T.; Data Curation: A.A.C., I.S.A., V.T., I.S.M., L.A., J.P.; Supervision: D.G.L., B.P., M.O., A.V., H.H., J.J.T., G.N., J.L.R.; Project Administration: G.N., J.L.R.; Funding Acquisition: G.N., J.L.R. J.L.R and G.N. wrote the manuscript with input from all other authors.

## Competing interests

The authors declare no competing interests.

## Notes

### Competing Interest Statement

The authors have declared no competing interest.

## References

1. Lopez-Rios, J. et al. Attenuated sensing of SHH by Ptch1 underlies evolution of bovine limbs. Nature 511, 46–51 (2014).

2. Cooper, K.L. et al. Patterning and post-patterning modes of evolutionary digit loss in mammals. Nature 511, 41–5 (2014).

3. Tissieres, V. et al. Gene Regulatory and Expression Differences between Mouse and Pig Limb Buds Provide Insights into the Evolutionary Emergence of Artiodactyl Traits. Cell Rep 31, 107490 (2020).

4. Saxena, A. et al. Interspecies transcriptomics identify genes that underlie disproportionate foot growth in jerboas. Curr Biol 32, 289–303 e6 (2022).

5. Schindler, M. et al. Comparative single-cell analyses reveal evolutionary repurposing of a conserved gene programme in bat wing development. Nat Ecol Evol 9, 1626–1642 (2025).

6. Carroll, S.B. Evo-devo and an expanding evolutionary synthesis: a genetic theory of morphological evolution. Cell 134, 25–36 (2008).

7. Osterwalder, M. et al. Enhancer redundancy provides phenotypic robustness in mammalian development. Nature 554, 239–243 (2018).

8. Cannavo, E. et al. Shadow Enhancers Are Pervasive Features of Developmental Regulatory Networks. Curr Biol 26, 38–51 (2016).

9. Abassah-Oppong, S. et al. A gene desert required for regulatory control of pleiotropic Shox2 expression and embryonic survival. Nat Commun 15, 8793 (2024).

10. Malkmus, J. et al. Spatial regulation by multiple Gremlin1 enhancers provides digit development with cis-regulatory robustness and evolutionary plasticity. Nat Commun 12, 5557 (2021).

11. Menke, D.B., Guenther, C. & Kingsley, D.M. Dual hindlimb control elements in the Tbx4 gene and region-specific control of bone size in vertebrate limbs. Development 135, 2543–53 (2008).

12. Sears, K.E. et al. Developmental basis of mammalian digit reduction: a case study in pigs. Evol Dev 13, 533–41 (2011).

13. Chen, Y. & Struhl, G. Dual roles for patched in sequestering and transducing hedgehog. Cell 87, 553–563 (1996).

14. Taipale, J., Cooper, M.K., Maiti, T. & Beachy, P.A. Patched acts catalytically to suppress the activity of Smoothened. Nature 418, 892–7 (2002).

15. Jeong, J. & McMahon, A.P. Growth and pattern of the mammalian neural tube are governed by partially overlapping feedback activities of the hedgehog antagonists patched 1 and Hhip1. Development 132, 143–54 (2005).

16. Marigo, V. & Tabin, C.J. Regulation of patched by sonic hedgehog in the developing neural tube. Proc Natl Acad Sci U S A 93, 9346–51 (1996).

17. Goodrich, L.V., Johnson, R.L., Milenkovic, L., McMahon, J.A. & Scott, M.P. Conservation of the *hedgehog/patched* signaling pathway from flies to mice: induction of a mouse *patched* gene by Hedgehog. Genes & Development 10, 301–312 (1996).

18. Vokes, S.A. et al. Genomic characterization of Gli-activator targets in sonic hedgehog-mediated neural patterning. Development 134, 1977–89 (2007).

19. Briscoe, J., Chen, Y., Jessell, T.M. & Struhl, G. A hedgehog-insensitive form of patched provides evidence for direct long-range morphogen activity of sonic hedgehog in the neural tube. Mol Cell 7, 1279–91 (2001).

20. Holtz, A.M. et al. Essential role for ligand-dependent feedback antagonism of vertebrate hedgehog signaling by PTCH1, PTCH2 and HHIP1 during neural patterning. Development 140, 3423–34 (2013).

21. Lorberbaum, D.S. et al. An ancient yet flexible cis-regulatory architecture allows localized Hedgehog tuning by patched/Ptch1. Elife 5(2016).

22. Desanlis, I., Paul, R. & Kmita, M. Transcriptional Trajectories in Mouse Limb Buds Reveal the Transition from Anterior-Posterior to Proximal-Distal Patterning at Early Limb Bud Stage. J Dev Biol 8(2020).

23. Markman, S. et al. A single-cell census of mouse limb development identifies complex spatiotemporal dynamics of skeleton formation. Dev Cell 58, 565–581 e4 (2023).

24. Palacio, V. et al. Single-cell profiling of penta- and tetradactyl mouse limb buds identifies mesenchymal progenitors controlling digit numbers and identities. Nat Commun 16, 1226 (2025).

25. Verheyden, J.M., Lewandoski, M., Deng, C., Harfe, B.D. & Sun, X. Conditional inactivation of Fgfr1 in mouse defines its role in limb bud establishment, outgrowth and digit patterning. Development 132, 4235–45 (2005).

26. Verheyden, J.M. & Sun, X. An Fgf/Gremlin inhibitory feedback loop triggers termination of limb bud outgrowth. Nature 454, 638–641. (2008).

27. Bravo Gonzalez-Blas, C., et al. SCENIC+: single-cell multiomic inference of enhancers and gene regulatory networks. Nat Methods 20, 1355–1367 (2023).

28. Mannion, B.J. et al. Uncovering hidden enhancers through unbiased in vivo testing. Nat Commun 16, 7313 (2025).

29. Hostikka, S.L. & Capecchi, M.R. The mouse Hoxc11 gene: genomic structure and expression pattern. Mech Dev 70, 133–45 (1998).

30. Nelson, C.E. et al. Analysis of *Hox* gene expression in the chick limb bud. Development 122, 1449–1466 (1996).

31. Riddle, R.D., Johnson, R.L., Laufer, E. & Tabin, C. Sonic hedgehog mediates the polarizing activity of the ZPA. Cell 75, 1401–16 (1993).

32. te Welscher, P., et al. Progression of Vertebrate Limb Development through SHH-Mediated Counteraction of GLI3. Science 298, 827–30 (2002).

33. Litingtung, Y., Dahn, R.D., Li, Y., Fallon, J.F. & Chiang, C. Shh and Gli3 are dispensable for limb skeleton formation but regulate digit number and identity. Nature 418, 979–83 (2002).

34. Zhu, J., Patel, R., Trofka, A., Harfe, B.D. & Mackem, S. Sonic hedgehog is not a limb morphogen but acts as a trigger to specify all digits in mice. Dev Cell 57, 2048–2062 e4 (2022).

35. Zhu, J. et al. Uncoupling Sonic hedgehog control of pattern and expansion of the developing limb bud. Dev Cell 14, 624–32 (2008).

36. Panman, L. et al. Differential regulation of gene expression in the digit forming area of the mouse limb bud by SHH and gremlin 1/FGF-mediated epithelial-mesenchymal signalling. Development 133, 3419–28 (2006).

37. Zuniga, A. et al. Conserved cis-regulatory regions in a large genomic landscape control SHH and BMP-regulated Gremlin1 expression in mouse limb buds. BMC Dev Biol 12, 23 (2012).

38. McLaughlin, M.T. et al. Frem1 activity is regulated by Sonic hedgehog signaling in the cranial neural crest mesenchyme during midfacial morphogenesis. Dev Dyn 252, 483–494 (2023).

39. Chuang, P.T. & McMahon, A.P. Vertebrate Hedgehog signalling modulated by induction of a Hedgehog-binding protein. Nature 397, 617–21 (1999).

40. Butterfield, N.C. et al. Patched 1 is a crucial determinant of asymmetry and digit number in the vertebrate limb. Development 136, 3515–24 (2009).

41. Lex, R.K. et al. GLI transcriptional repression regulates tissue-specific enhancer activity in response to Hedgehog signaling. Elife 9(2020).

42. Sheth, R. et al. Distal Limb Patterning Requires Modulation of cis-Regulatory Activities by HOX13. Cell Rep 17, 2913–2926 (2016).

43. Goodrich, L.V., Milenkovic, L., Higgins, K.M. & Scott, M.P. Altered neural cell fates and medulloblastoma in mouse patched mutants. Science 277, 1109–13. (1997).

44. Artus, J. & Hadjantonakis, A.K. Generation of chimeras by aggregation of embryonic stem cells with diploid or tetraploid mouse embryos. Methods Mol Biol 693, 37–56 (2011).

45. Visel, A., Minovitsky, S., Dubchak, I. & Pennacchio, L.A. VISTA Enhancer Browser--a database of tissue-specific human enhancers. Nucleic Acids Res 35, D88–92 (2007).

46. Kosicki, M. et al. VISTA Enhancer browser: an updated database of tissue-specific developmental enhancers. Nucleic Acids Res 53, D324–D330 (2025).

47. Schwartzman, O. et al. UMI-4C for quantitative and targeted chromosomal contact profiling. Nat Methods 13, 685–91 (2016).

48. Stuart, T., Srivastava, A., Madad, S., Lareau, C.A. & Satija, R. Single-cell chromatin state analysis with Signac. Nat Methods 18, 1333–1341 (2021).

49. Cooper, K.L. The case against simplistic genetic explanations of evolution. Development 151(2024).

50. Benazet, J.D. et al. A self-regulatory system of interlinked signaling feedback loops controls mouse limb patterning. Science 323, 1050–3 (2009).

51. Zuniga, A., Haramis, A.P., McMahon, A.P. & Zeller, R. Signal relay by BMP antagonism controls the SHH/FGF4 feedback loop in vertebrate limb buds. Nature 401, 598–602. (1999).

52. Sharpe, J., Nonchev, S., Gould, A., Whiting, J. & Krumlauf, R. Selectivity, sharing and competitive interactions in the regulation of Hoxb genes. EMBO J 17, 1788–98 (1998).

53. Montavon, T. et al. A regulatory archipelago controls Hox genes transcription in digits. Cell 147, 1132–45 (2011).

54. Berlivet, S. et al. Clustering of tissue-specific sub-TADs accompanies the regulation of HoxA genes in developing limbs. PLoS Genet 9, e1004018 (2013).

55. Andrey, G. et al. A switch between topological domains underlies HoxD genes collinearity in mouse limbs. Science 340, 1234167 (2013).

56. Rancourt, D.E., Tsuzuki, T. & Capecchi, M.R. Genetic interaction between hoxb-5 and hoxb-6 is revealed by nonallelic noncomplementation. Genes Dev 9, 108–22 (1995).

57. Medina-Martinez, O., Bradley, A. & Ramirez-Solis, R. A large targeted deletion of Hoxb1-Hoxb9 produces a series of single-segment anterior homeotic transformations. Dev Biol 222, 71–83 (2000).

58. Xu, B. et al. Hox5 interacts with Plzf to restrict Shh expression in the developing forelimb. Proc Natl Acad Sci U S A 110, 19438–43 (2013).

59. Martinez-Abadias, N. et al. Quantification of gene expression patterns to reveal the origins of abnormal morphogenesis. Elife 7(2018).

60. Caskenette, D. et al. Global deletion of Panx3 produces multiple phenotypic effects in mouse humeri and femora. J Anat 228, 746–56 (2016).

61. Wellik, D.M. & Capecchi, M.R. Hox10 and Hox11 genes are required to globally pattern the mammalian skeleton. Science 301, 363–7 (2003).

62. Prothero, D.R., Foss, S.E. & (eds). The evolution of artiodactyls. The evolution of artiodactyls., i-xii, 1–367 (2007).

63. Hornblad, A., Bastide, S., Langenfeld, K., Langa, F. & Spitz, F. Dissection of the Fgf8 regulatory landscape by in vivo CRISPR-editing reveals extensive intra- and inter-enhancer redundancy. Nat Commun 12, 439 (2021).

64. Iliadou, M., Haro, E. & Rada-Iglesias, A. Mechanisms and functional implications of long-range enhancer-dependent gene regulation. Nat Genet 58, 2104–2117 (2026).

65. Inoue, F., Kreimer, A., Ashuach, T., Ahituv, N. & Yosef, N. Identification and Massively Parallel Characterization of Regulatory Elements Driving Neural Induction. Cell Stem Cell 25, 713–727 e10 (2019).

66. Rouco, R. et al. Temporal constraints on enhancer usage shape the regulation of limb gene transcription. Nat Commun 17, 5 (2026).

67. Lopez-Rios, J. et al. GLI3 constrains digit number by controlling both progenitor proliferation and BMP-dependent exit to chondrogenesis. Dev Cell 22, 837–48 (2012).

68. Love, M.I., Huber, W. & Anders, S. Moderated estimation of fold change and dispersion for RNA-seq data with DESeq2. Genome Biol 15, 550 (2014).

69. Ramos-Rodriguez, M., Subirana-Granes, M. & Pasquali, L. UMI4Cats: an R package to analyze chromatin contact profiles obtained by UMI-4C. Bioinformatics 37, 4240–4242 (2021).

70. Harfe, B.D. et al. Evidence for an expansion-based temporal Shh gradient in specifying vertebrate digit identities. Cell 118, 517–28 (2004).

71. George, S.H. et al. Developmental and adult phenotyping directly from mutant embryonic stem cells. Proc Natl Acad Sci U S A 104, 4455–60 (2007).

72. Kraft, K. et al. Deletions, Inversions, Duplications: Engineering of Structural Variants using CRISPR/Cas in Mice. Cell Rep 10, 833–839 (2015).

73. Ran, F.A. et al. Genome engineering using the CRISPR-Cas9 system. Nat Protoc 8, 2281–308 (2013).

74. Modzelewski, A.J. et al. Efficient mouse genome engineering by CRISPR-EZ technology. Nat Protoc 13, 1253–1274 (2018).

75. Stuart, T. et al. Comprehensive Integration of Single-Cell Data. Cell 177, 1888–1902 e21 (2019).

76. Korsunsky, I. et al. Fast, sensitive and accurate integration of single-cell data with Harmony. Nat Methods 16, 1289–1296 (2019).

77. Rueda, A.D. et al. The cellular landscape of the endochondral bone during the transition to extrauterine life. Immunol Cell Biol 102, 131–148 (2024).

78. Zhang, Y. et al. Model-based analysis of ChIP-Seq (MACS). Genome Biol 9, R137 (2008).

79. Zhao, Y. et al. A compendium and comparative epigenomics analysis of cis-regulatory elements in the pig genome. Nat Commun 12, 2217 (2021).

80. Delgado, I. et al. Control of mouse limb initiation and antero-posterior patterning by Meis transcription factors. Nat Commun 12, 3086 (2021).

81. Gene Ontology, C. The Gene Ontology resource: enriching a GOld mine. Nucleic Acids Res 49, D325–D334 (2021).

82. Wu, T. et al. clusterProfiler 4.0: A universal enrichment tool for interpreting omics data. Innovation (Camb) 2, 100141 (2021).

83. Boyle, E.I. et al. GO::TermFinder--open source software for accessing Gene Ontology information and finding significantly enriched Gene Ontology terms associated with a list of genes. Bioinformatics 20, 3710–5 (2004).

84. Corces, M.R. et al. The chromatin accessibility landscape of primary human cancers. Science 362(2018).

85. Mimno, D., Wallach, H., Talley, E., Leenders, M. & McCallum, A. Optimizing Semantic Coherence in Topic Models. 262–272 (Association for Computational Linguistics, Edinburgh, Scotland, UK., 2011).

86. Griffiths, T.L. & Steyvers, M. Finding scientific topics. Proc Natl Acad Sci U S A 101 Suppl 1, 5228–35 (2004).

87. Frith, M.C., Li, M.C. & Weng, Z. Cluster-Buster: Finding dense clusters of motifs in DNA sequences. Nucleic Acids Res 31, 3666–8 (2003).

88. Bastian, M., Heymann, S. & Jacomy, M. Gephi: An Open Source Software for Exploring and Manipulating Networks. Proceedings of the International AAAI Conference on Web and Social Media 3, 361–362 (2009).

89. Andrey, G. et al. Characterization of hundreds of regulatory landscapes in developing limbs reveals two regimes of chromatin folding. Genome Res 27, 223–233 (2017).

90. Sospedra-Arrufat, I. et al. Epigenomic signatures of cis-regulatory elements in the developing mouse and pig forelimb. Sci Data 12, 598 (2025).

91. Kraft, K. et al. Serial genomic inversions induce tissue-specific architectural stripes, gene misexpression and congenital malformations. Nat Cell Biol 21, 305–310 (2019).

