## Supplementary material for "Network inference at single-cell resolution reveals morphoregulatory divergence and multilayered buffering in limb evolution": Suplementary Figures

### Figure S1

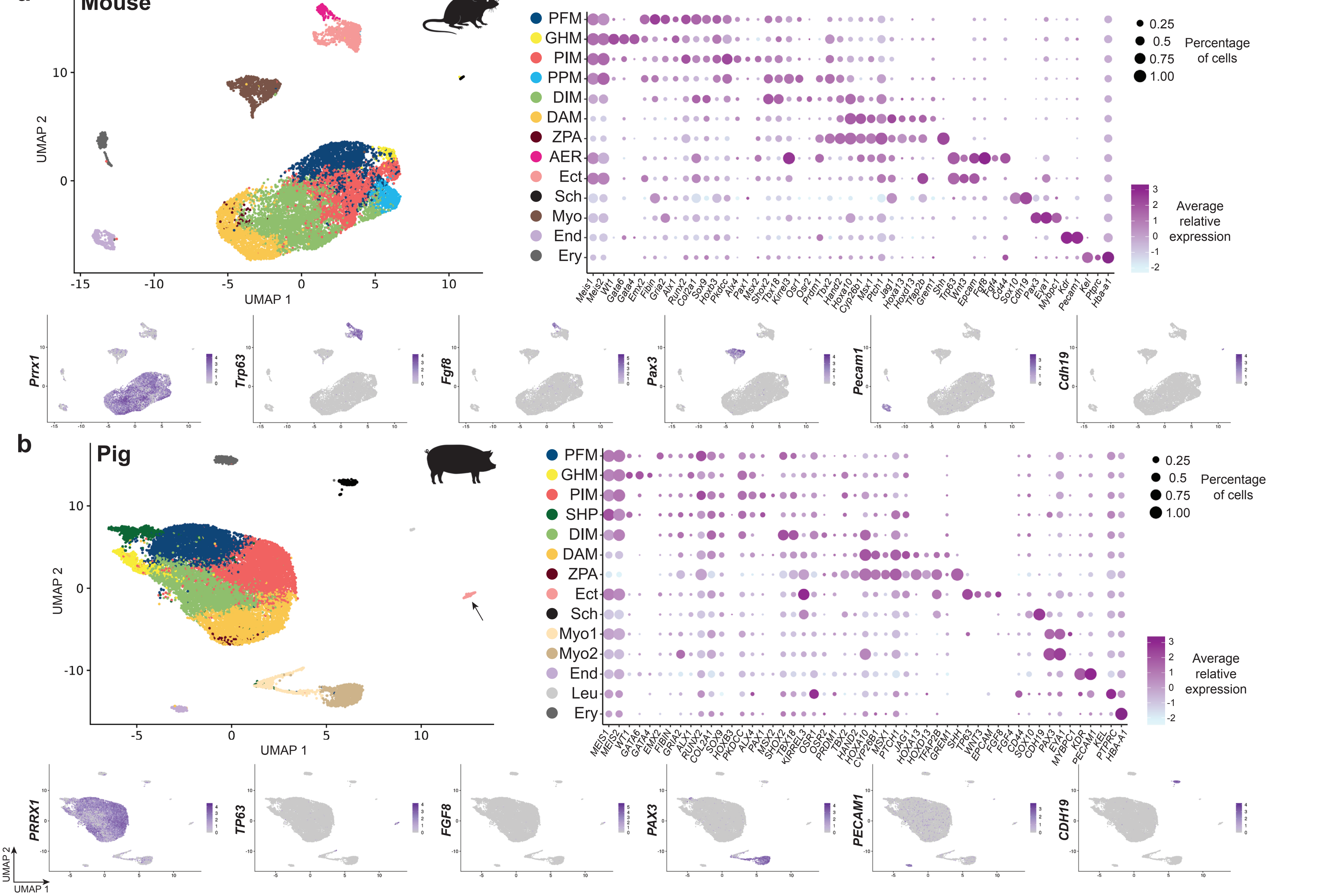

a
Mouse
UMAP 2
UMAP 1
PFM
GHM
PIM
PPM
DIM
DAM
ZPA
AER
Ect
Sch
Myo
End
Ery
Meis1 Meis2 Wt1 Gata6 Gata4 Emx2 Fibin Gria2 Alx1 Runx2 Col2a1 Sox9 Hoxb3 Pkdcc Alx4 Pax1 Msx2 Shox2 Tbx18 Kirrel3 Osr1 Osr2 Prdm1 Tbx2 Hand2 Hoxa10 Cyp26b1 Msx1 Ptch1 Jag1 Hoxa13 Hoxd13 Tfap2b Grem1 Shh Trp63 Wnt3 Epcam Fgf8 Fgf4 Cd44 Sox10 Cdh19 Pax3 Eya1 Mybpc1 Kdr Pecam1 Kel Ptprc Hba-a1
Percentage of cells
0.25
0.5
0.75
1.00
Average relative expression
Prrx1
Trp63
Fgf8
Pax3
Pecam1
Cdh19
b
Pig
PFM
GHM
PIM
SHP
DIM
DAM
ZPA
Ect
Sch
Myo1
Myo2
End
Leu
Ery
MEIS1 MEIS2 WT1 GATA6 GATA4 EMX2 FIBIN GRIA2 ALX1 RUNX2 COL2A1 SOX9 HOXB3 PKDCC ALX4 PAX1 MSX2 SHOX2 TBX18 KIRREL3 OSR1 OSR2 PRDM1 TBX2 HAND2 HOXA10 CYP26B1 MSX1 PTCH1 JAG1 HOXA13 HOXD13 TFAP2B GREM1 SHH TP63 WNT3 EPCAM FGF8 FGF4 CD44 SOX10 CDH19 PAX3 EYA1 MYBPC1 KDR PECAM1 KEL PTPRC HBA-A1
Percentage of cells
0.25
0.5
0.75
1.00
Average relative expression
PRRX1
TP63
FGF8
PAX3
PECAM1
CDH19
UMAP 2
UMAP 1

### Figure S2

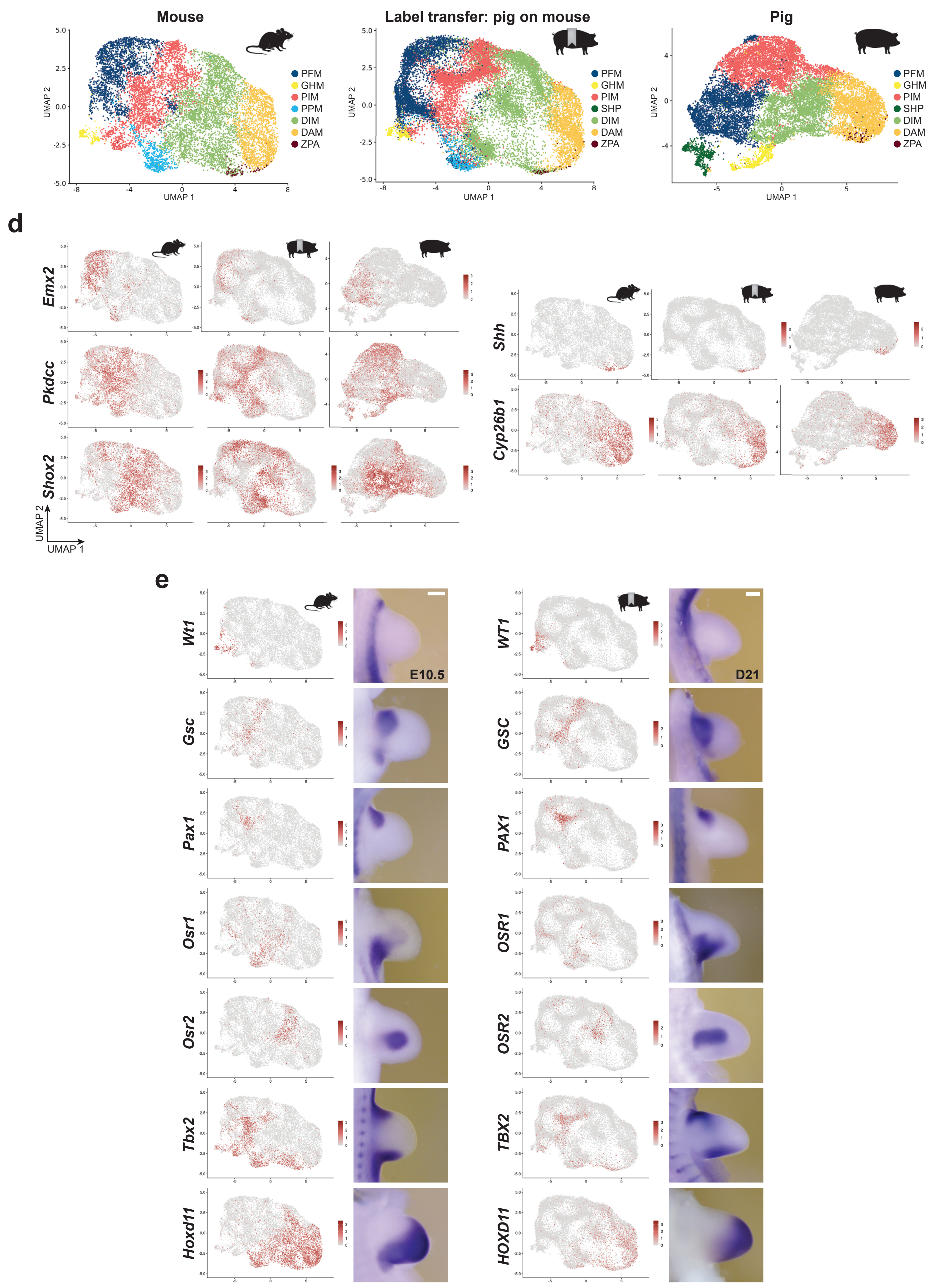

a
b
c
Mouse
Label transfer: pig on mouse
Pig
PFM
GHM
PIM
PPM
SHP
DIM
DAM
ZPA
UMAP 1
UMAP 2
d
Emx2
Pkdcc
Shox2
Shh
Cyp26b1
e
Wt1
Gsc
Pax1
Osr1
Osr2
Tbx2
Hoxd11
WT1
GSC
PAX1
OSR1
OSR2
TBX2
HOXD11
E10.5
D21

### Figure S3

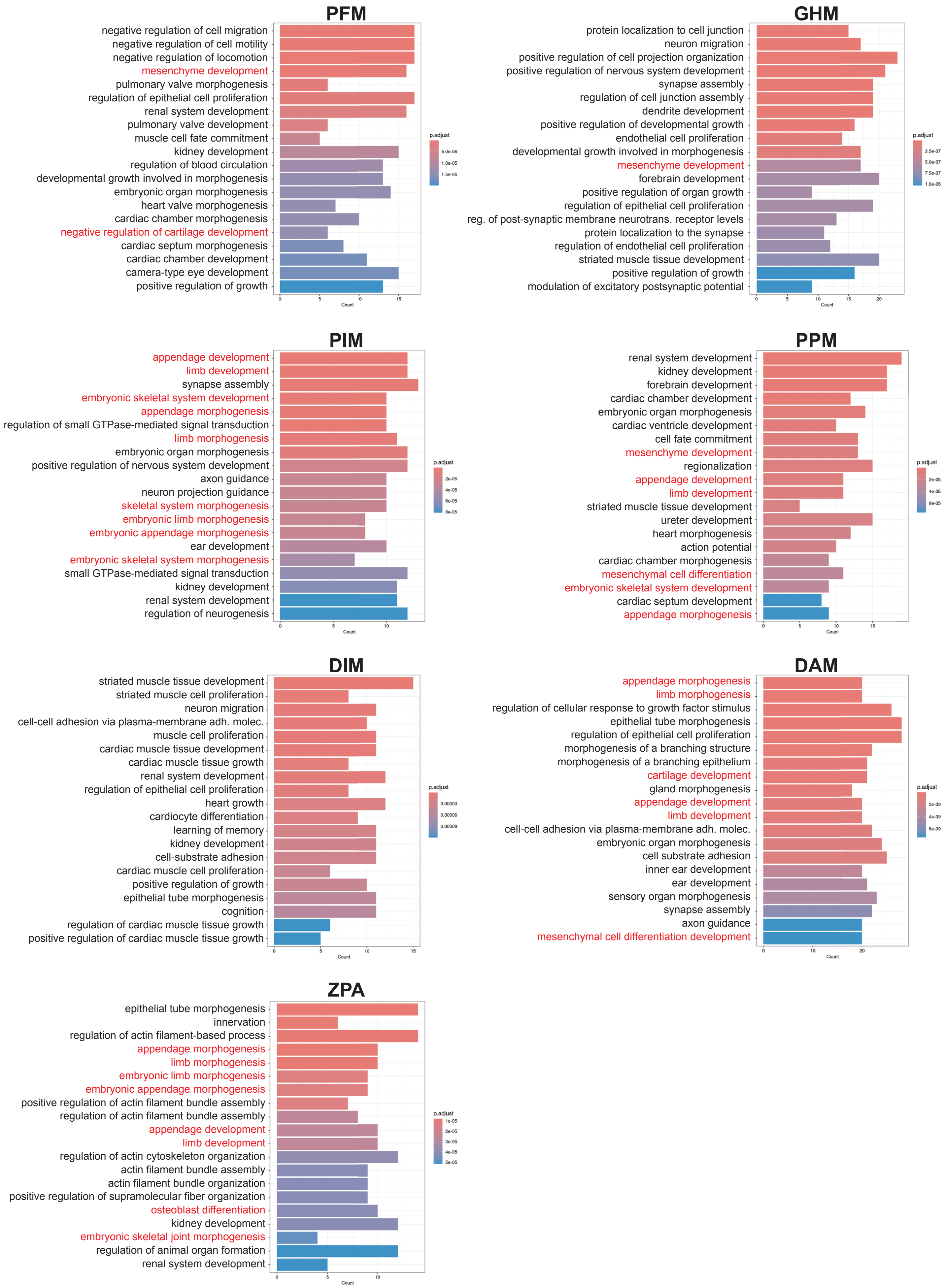
PFM
negative regulation of cell migration
negative regulation of cell motility
negative regulation of locomotion
mesenchyme development
pulmonary valve morphogenesis
regulation of epithelial cell proliferation
renal system development
pulmonary valve development
muscle cell fate commitment
kidney development
regulation of blood circulation
developmental growth involved in morphogenesis
embryonic organ morphogenesis
heart valve morphogenesis
cardiac chamber morphogenesis
negative regulation of cartilage development
cardiac septum morphogenesis
cardiac chamber development
camera-type eye development
positive regulation of growth
Count
p.adjust
GHM
protein localization to cell junction
neuron migration
positive regulation of cell projection organization
positive regulation of nervous system development
synapse assembly
regulation of cell junction assembly
dendrite development
positive regulation of developmental growth
endothelial cell proliferation
developmental growth involved in morphogenesis
mesenchyme development
forebrain development
positive regulation of organ growth
regulation of epithelial cell proliferation
reg. of post-synaptic membrane neurotrans. receptor levels
protein localization to the synapse
regulation of endothelial cell proliferation
striated muscle tissue development
positive regulation of growth
modulation of excitatory postsynaptic potential
Count
p.adjust
PIM
appendage development
limb development
synapse assembly
embryonic skeletal system development
appendage morphogenesis
regulation of small GTPase-mediated signal transduction
limb morphogenesis
embryonic organ morphogenesis
positive regulation of nervous system development
axon guidance
neuron projection guidance
skeletal system morphogenesis
embryonic limb morphogenesis
embryonic appendage morphogenesis
ear development
embryonic skeletal system morphogenesis
small GTPase-mediated signal transduction
kidney development
renal system development
regulation of neurogenesis
Count
p.adjust
PPM
renal system development
kidney development
forebrain development
cardiac chamber development
embryonic organ morphogenesis
cardiac ventricle development
cell fate commitment
mesenchyme development
regionalization
appendage development
limb development
striated muscle tissue development
ureter development
heart morphogenesis
action potential
cardiac chamber morphogenesis
mesenchymal cell differentiation
embryonic skeletal system development
cardiac septum development
appendage morphogenesis
Count
p.adjust
DIM
striated muscle tissue development
striated muscle cell proliferation
neuron migration
cell-cell adhesion via plasma-membrane adh. molec.
muscle cell proliferation
cardiac muscle tissue development
cardiac muscle tissue growth
renal system development
regulation of epithelial cell proliferation
heart growth
cardiocyte differentiation
learning of memory
kidney development
cell-substrate adhesion
cardiac muscle cell proliferation
positive regulation of growth
epithelial tube morphogenesis
cognition
regulation of cardiac muscle tissue growth
positive regulation of cardiac muscle tissue growth
Count
p.adjust
DAM
appendage morphogenesis
limb morphogenesis
regulation of cellular response to growth factor stimulus
epithelial tube morphogenesis
regulation of epithelial cell proliferation
morphogenesis of a branching structure
morphogenesis of a branching epithelium
cartilage development
gland morphogenesis
appendage development
limb development
cell-cell adhesion via plasma-membrane adh. molec.
embryonic organ morphogenesis
cell substrate adhesion
inner ear development
ear development
sensory organ morphogenesis
synapse assembly
axon guidance
mesenchymal cell differentiation development
Count
p.adjust
ZPA
epithelial tube morphogenesis
innervation
regulation of actin filament-based process
appendage morphogenesis
limb morphogenesis
embryonic limb morphogenesis
embryonic appendage morphogenesis
positive regulation of actin filament bundle assembly
regulation of actin filament bundle assembly
appendage development
limb development
regulation of actin cytoskeleton organization
actin filament bundle assembly
actin filament bundle organization
positive regulation of supramolecular fiber organization
osteoblast differentiation
kidney development
embryonic skeletal joint morphogenesis
regulation of animal organ formation
renal system development
Count
p.adjust

### Figure S4

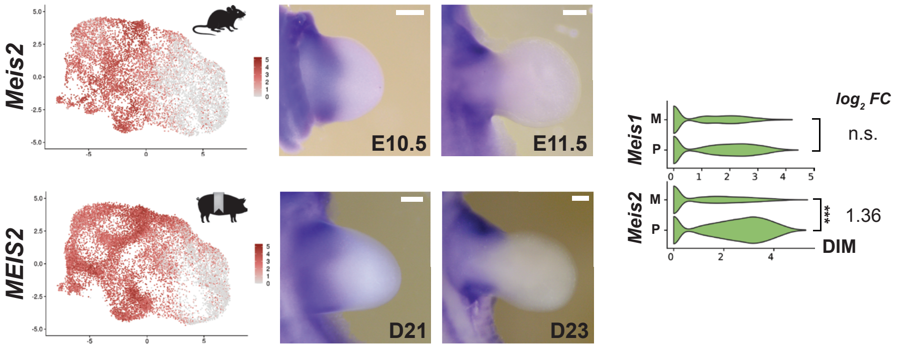

Meis2
MEIS2
E10.5
E11.5
D21
D23
log2 FC
Meis1
Meis2
M
P
n.s.
***
1.36
DIM

### Figure S5

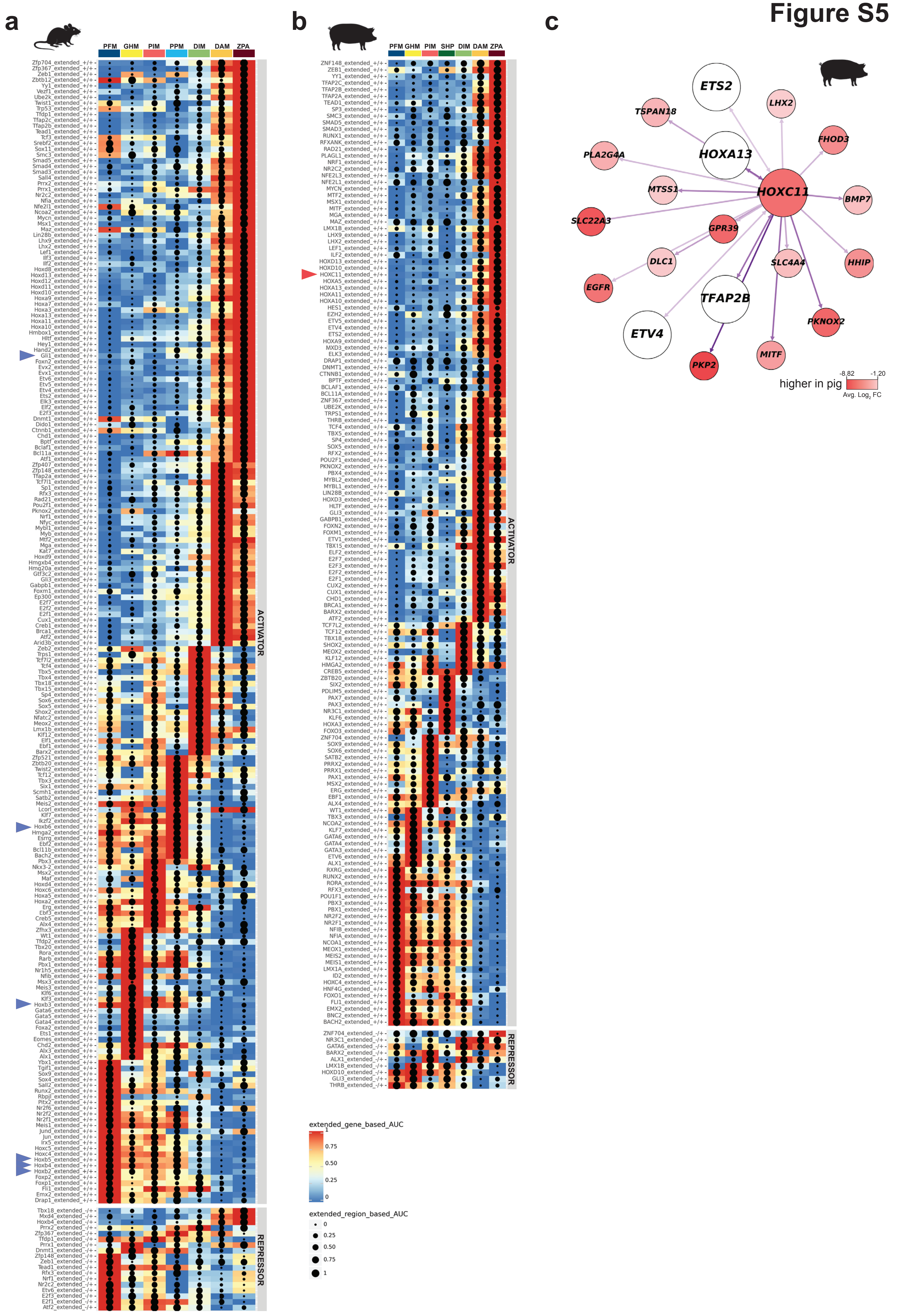

a
PFM
GHM
PIM
PPM
DIM
DAM
ZPA
ACTIVATOR
REPRESSOR
b
PFM
GHM
PIM
SHP
DIM
DAM
ZPA
ACTIVATOR
REPRESSOR
extended_gene_based_AUC
1
0.75
0.50
0.25
0
extended_region_based_AUC
0
0.25
0.50
0.75
1
c
ETS2
LHX2
TSPAN18
FHOD3
PLA2G4A
HOXA13
MTSS1
HOXC11
BMP7
SLC22A3
GPR39
DLC1
SLC4A4
HHIP
EGFR
TFAP2B
PKNOX2
ETV4
MITF
PKP2
higher in pig
-8.82
-1.20
Avg. Log2 FC

Figure S6

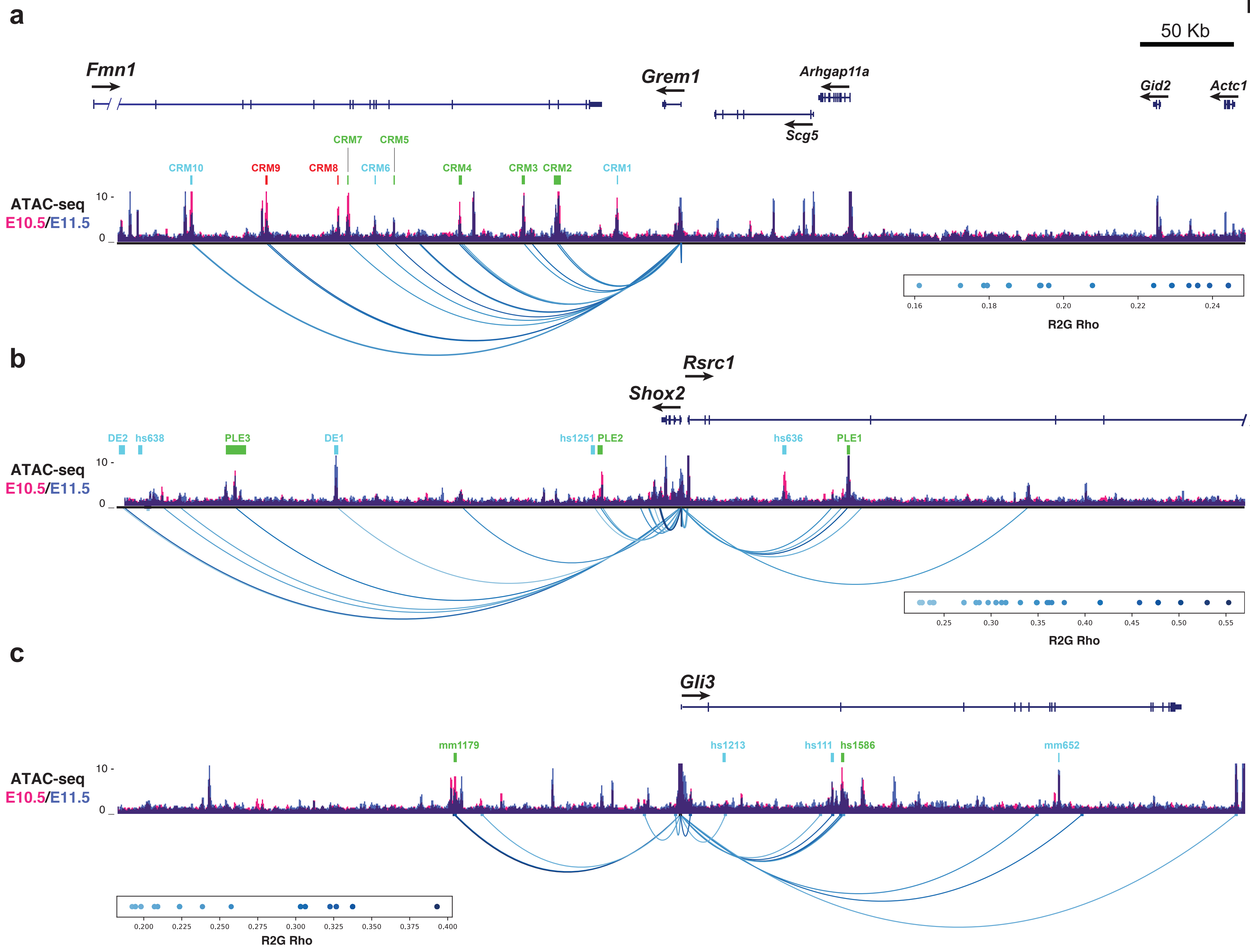
a
50 Kb
Fmn1
Grem1
Arhgap11a
Gid2
Actc1
Scg5
CRM7
CRM5
CRM10
CRM9
CRM8
CRM6
CRM4
CRM3
CRM2
CRM1
ATAC-seq
E10.5/E11.5
10 -
0 _
0.16
0.18
0.20
0.22
0.24
R2G Rho
b
Rsrc1
Shox2
DE2
hs638
PLE3
DE1
hs1251
PLE2
hs636
PLE1
ATAC-seq
E10.5/E11.5
10 -
0 _
0.25
0.30
0.35
0.40
0.45
0.50
0.55
R2G Rho
c
Gli3
mm1179
hs1213
hs111
hs1586
mm652
ATAC-seq
E10.5/E11.5
10 -
0 _
0.200
0.225
0.250
0.275
0.300
0.325
0.350
0.375
0.400
R2G Rho

### Figure S7

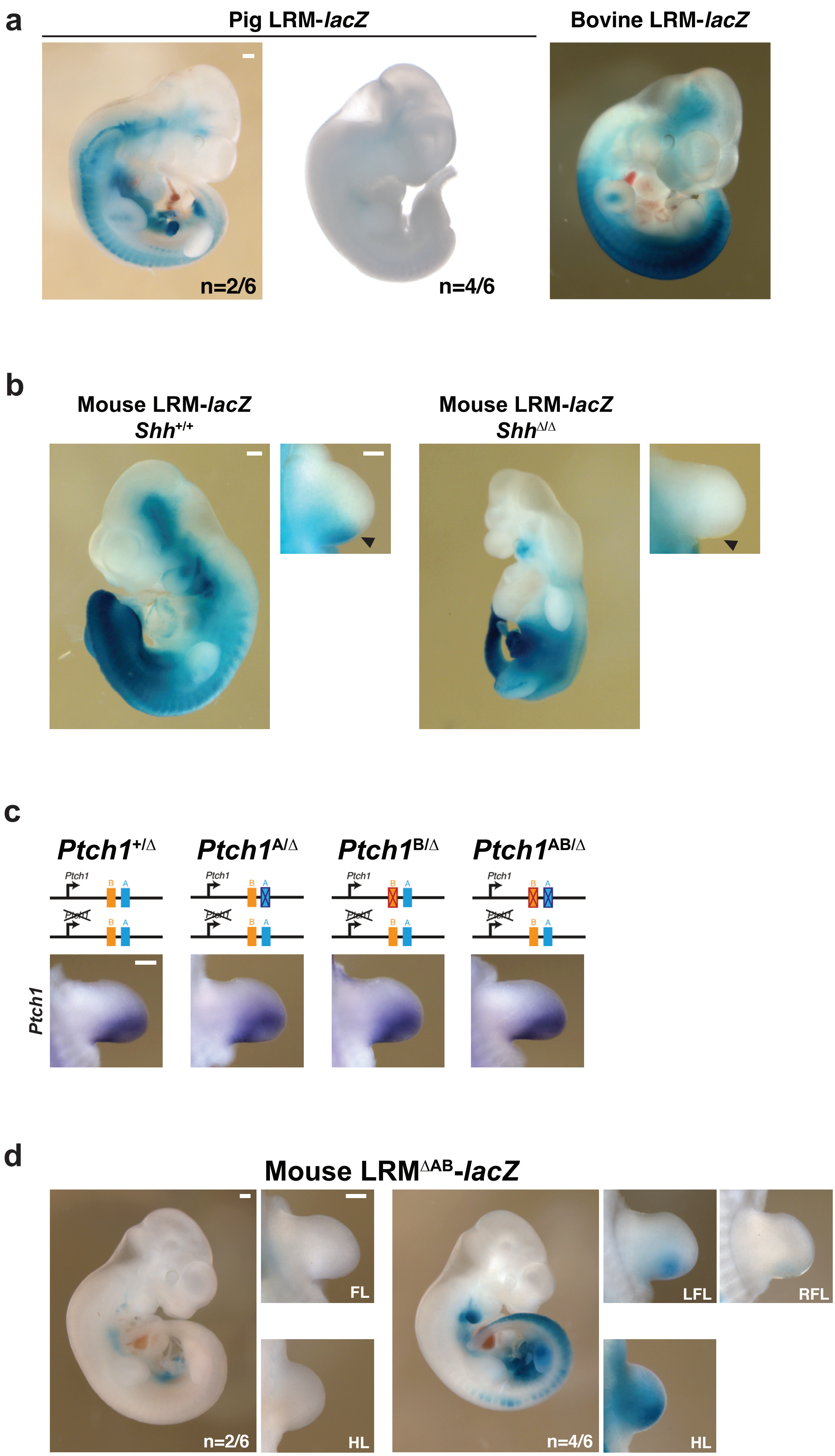
a
Pig LRM-lacZ
Bovine LRM-lacZ
n=2/6
n=4/6
b
Mouse LRM-lacZ
Shh+/+
Mouse LRM-lacZ
ShhΔ/Δ
c
Ptch1+/Δ
Ptch1A/Δ
Ptch1B/Δ
Ptch1AB/Δ
Ptch1
B
A
Ptch1
d
Mouse LRMΔAB-lacZ
FL
HL
LFL
RFL
HL
n=2/6
n=4/6

### Figure S8

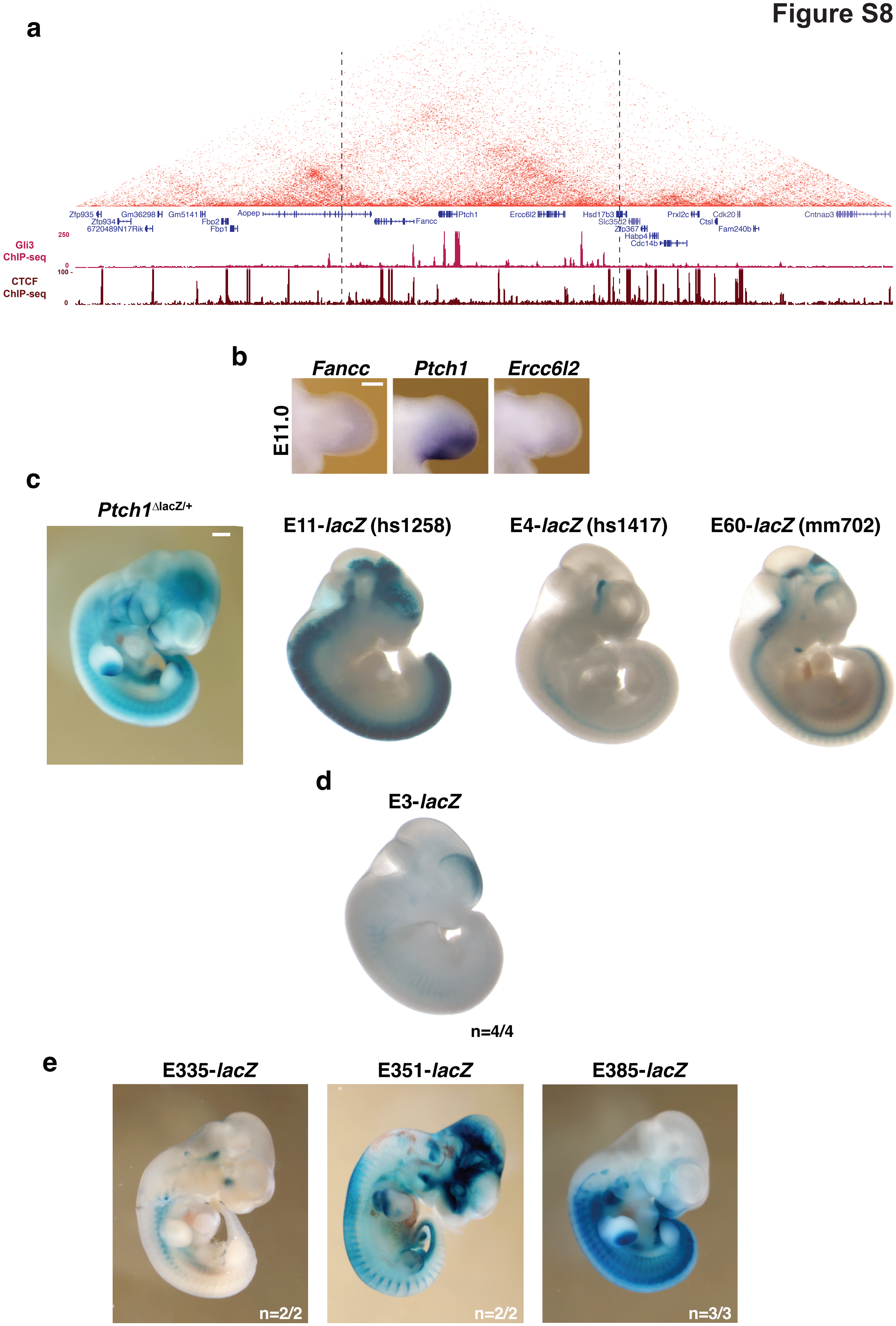

a
Zfp935
Zfp934
6720489N17Rik
Gm36298
Gm5141
Fbp2
Fbp1
Aopep
Fancc
Ptch1
Ercc6l2
Hsd17b3
Slc35d2
Zfp367
Habp4
Cdc14b
Prxl2c
Ctsl
Cdk20
Fam240b
Cntnap3
250
0
100 -
0
Gli3
ChIP-seq
CTCF
ChIP-seq
b
Fancc
Ptch1
Ercc6l2
E11.0
c
Ptch1ΔlacZ/+
E11-lacZ (hs1258)
E4-lacZ (hs1417)
E60-lacZ (mm702)
d
E3-lacZ
n=4/4
e
E335-lacZ
E351-lacZ
E385-lacZ
n=2/2
n=2/2
n=3/3

### Figure S9

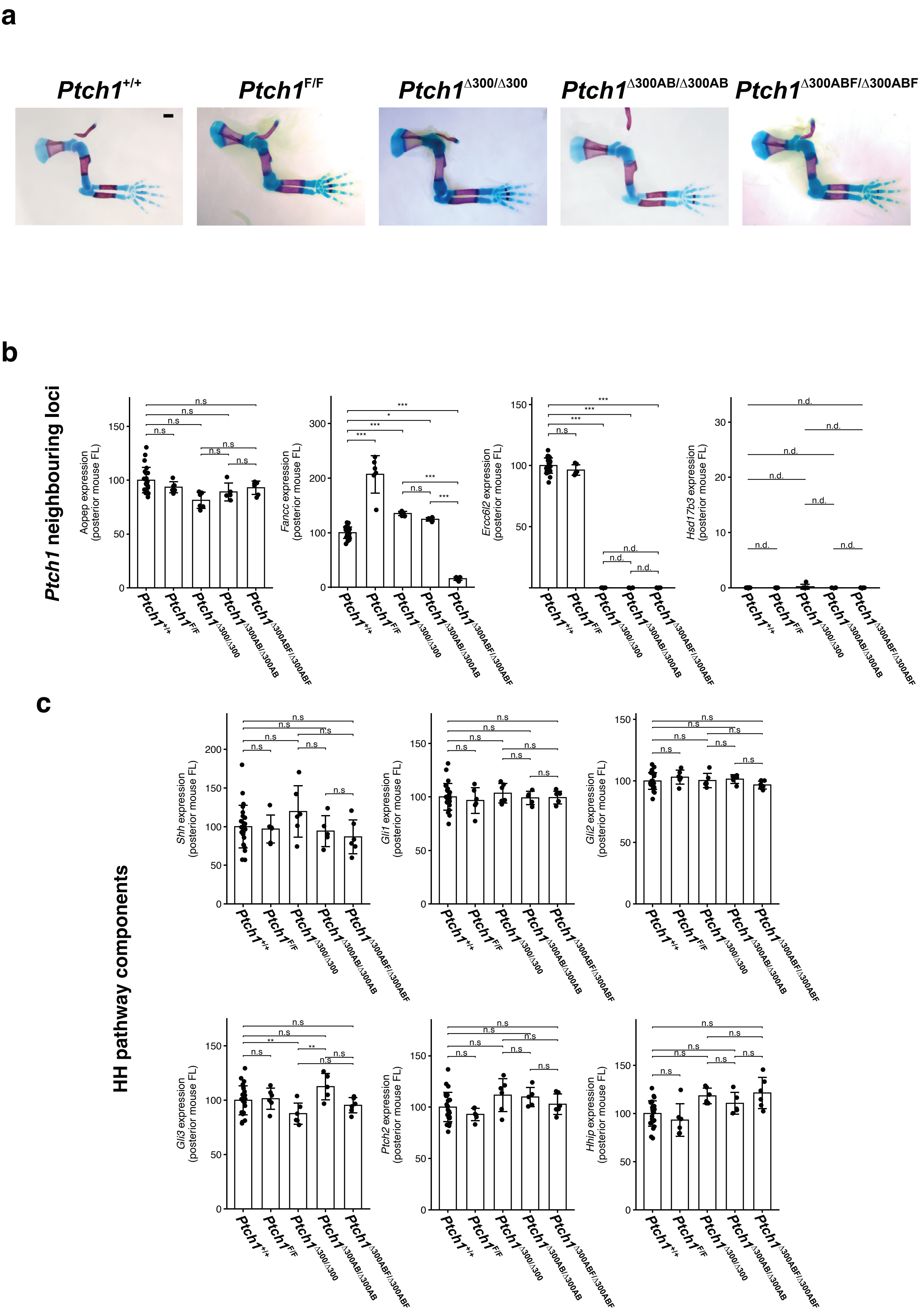
a
$Ptch1^{+/+}$
$Ptch1^{F/F}$
$Ptch1^{\Delta300/\Delta300}$
$Ptch1^{\Delta300AB/\Delta300AB}$
$Ptch1^{\Delta300ABF/\Delta300ABF}$
b
Ptch1 neighbouring loci
Aopep expression (posterior mouse FL)
Fancc expression (posterior mouse FL)
Ercc6l2 expression (posterior mouse FL)
Hsd17b3 expression (posterior mouse FL)
n.s
***
*
n.d.
c
HH pathway components
Shh expression (posterior mouse FL)
Gli1 expression (posterior mouse FL)
Gli2 expression (posterior mouse FL)
Gli3 expression (posterior mouse FL)
Ptch2 expression (posterior mouse FL)
Hhip expression (posterior mouse FL)
**
